# Benchmarking temporal causal discovery for fully and partially observed biochemical kinetic models

**DOI:** 10.64898/2026.11.12.766185

**Authors:** Holly Chambers, Hervé Isambert, Vahid Shahrezaei, Barbara Bravi

## Abstract

Accurately reconstructing networks of biochemical reactions is a crucial yet unsolved problem for leveraging advances in experimental techniques that track intracellular dynamics in a time-resolved fashion. This task is further challenged by the fact that we typically observe only part of the system, while many variables shaping the dynamics remain unobserved. Causal discovery methods can infer causal relations from purely observational data and account for unobserved variables. They therefore hold promise for yielding faithful, mechanistically interpretable representations of underlying biochemical interactions, but their potential in this context remains underexplored.

In this work, we provide a systematic benchmark of state-of-the-art temporal causal discovery methods (tMIIC, PCMCI+/LPCMCI and SVAR-FCI) on time-series data from simulations of biochemical kinetics of mass action and Michaelis-Menten type modelled as systems of nonlinear stochastic differential equations, including an experimentally calibrated model of the epidermal growth factor receptor signalling network. By selectively omitting data, we consider the problem of reconstructing these networks in the presence of both latent confounders and unobserved species participating in reactions. We find that tMIIC and PCMCI+/LPCMCI achieve the highest overall prediction quality, with tMIIC able to handle larger samples within the same runtime budget. We identify, as the main condition for optimal performance of causal discovery, the consistency between data-sampling regimes (sampling intervals, trajectory length) and the typical timescales of the system. We establish that causal discovery can detect latent confounders via bidirected edges and unobserved species via time-delayed edges, locating hidden effects and encoding information on their typical timescales, provided that the data sampling guarantees adequate temporal resolution. As such, our results can guide the choice of hyperparameters in causal discovery algorithms and of data-sampling regimes to optimize network-reconstruction performance.

**Author summary:** Systems of intracellular biochemical reactions are highly complex. Inferring, from observational data, network representations of these systems, with nodes representing biochemical species and edges representing their reactions, is of fundamental importance for quantitatively characterizing their function and the effects of dysregulation, yet it remains challenging. A further difficulty is that biochemical systems often involve components that cannot be directly measured, although we would like to know if and how these ‘hidden nodes’ influence what we observe. Causal discovery methods, i.e. methods that can learn networks of causal relationships, have the potential to bring substantial improvements, one of their strengths being their mechanisms for detecting hidden-node effects from data.

In this work, we deploy and benchmark causal discovery to learn causal networks from time-series data generated by models of fully or partially observed stochastic biochemical kinetics, comparing state-of-the-art methods and identifying the sampling-regime characteristics that lead to optimal performance relative to the typical timescales of the kinetics. Our results demonstrate the ability of these methods to reconstruct the original biochemical networks from which the data were generated and to detect hidden nodes, showcasing the feasibility and usefulness of causal discovery as part of data-driven mathematical-modelling pipelines for biochemical reaction systems.

## Introduction

The problem of reconstructing biochemical kinetic networks from observational data is at the forefront of computational molecular biology, motivated by recent advancements in experimental techniques to monitor gene expression and biochemical signalling, often at the single-cell level and with temporal resolution [1]. On the one hand, live-cell imaging with high-resolution microscopy enables the observation of cellular dynamics, from organelle behaviour and vesicle trafficking [2] to microbial life cycle stages [3]. On the other hand, modern single-cell sequencing technologies with scRNA-seq methods, despite yielding only snapshot data, are amenable to the inference of pseudotime gene expression trajectories [4] and have been extended to sequential transcriptomic profiling over time (e.g., live-seq [5]). Accurate network reconstruction in these contexts can provide insight into how cellular components interact, and how interventions alter their behaviour, providing a foundation for predictive, mechanistic models.

Reconstruction of the true underlying structure of complex intracellular biochemical networks remains particularly challenging for several reasons, first because it is far from straightforward to disentangle the effect of direct, biophysical molecular interactions from indirect correlations [6]. But also because, in practice, measurements are inevitably sparse and thus provide only *partial* observations: subnetworks embedded in a larger, unobserved whole which we refer to as ‘bulk’. For instance, the challenge inherent to the behaviour of observations being influenced by components that are not directly observable has been highlighted in work on complex systems in biology [7, 8], and on live-cell imaging of cellular features through time [9].

When bulk species are marginalized out, their influence manifests as non-Markovian memory effects in the reduced equations governing the observed variables, as established by theoretical frameworks for coarse-grained stochastic dynamics [10–13]. These memory effects act on boundary species (observed variables with influence from the bulk), with timescales that reflect the rates of the hidden nodes. Recent work has used physics-inspired machine learning to identify network boundaries distinguishing them from interior nodes precisely by detecting the non-Markovian effects that are unique to the dynamics of these boundary species [14].

Causal discovery methods provide a framework for inferring causal networks from observational data, both of static and dynamical nature. Since they are designed to tackle the problem of disentangling direct causation from indirect correlations, with some techniques capable of indicating the presence of hidden variables, they are ideally suited to drive the next performance improvements at biochemical network reconstruction. There are two main classes of causal discovery methods: score-based methods search the space of potential network structures assigning some numerical score representing how well a fit to the data has been made [15–17], while constraint-based methods use statistical tests of conditional independence to identify putative causally related variables, removing and orienting edges according to the statistical dependencies observed in the data [18, 19]. For time series data, constraint-based methods operate on time-unfolded causal networks (described in Box 1), and several extensions have been developed to handle latent confounders and contemporaneous effects [20–22] . Here, we focus on three state-of-the-art temporal (and its latent variable extension LPCMCI [21]), constraint-based algorithms – tMIIC [23], PCMCI+ and SVAR-FCI [24].

Early approaches to network inference of gene-gene interactions and gene regulatory networks (GRNs) relied on perturbation-based, non-causal methods, for example [25]. Subsequent work includes explicitly causal approaches, for example in work using static variants of causal discovery to reconstruct GRNs [26–28]. For time series gene expression data, Granger causality [29] has been a dominant framework for inferring relationships [30–33], though without reference to a graphical model, and with no mechanism for detecting hidden variables.

In light of these promising applications to complex biological dynamics, recent work has focussed on benchmarking causal discovery methods. In [34], twenty static causal discovery methods are tested on data from the UK Biobank project with the aim of identifying the causal effect of particulate matter on incidence of cardiovascular disease. A benchmarking platform for temporal causal discovery, CausalDynamics was introduced by [35], including scenarios with unobserved confounders. However, a systematic benchmark of temporal causal discovery specifically in the context of biochemical kinetics (with nonlinearities and multiplicative noise) is currently lacking from the literature. In this context, in the case of partially observed data, a relevant, still unaddressed scenario is the one where the unobserved are not simply pure-noise latent confounders but biochemical species participating in the reactions of the network. A natural question in this scenario is to what extent and under which conditions causal discovery can detect the presence of such species and infer information about their dynamical effect onto the observed network species.

Here, we consider three state-of-the-art causal discovery algorithms (tMIIC, PCMCI+/LPCMCI and SVAR-FCI) that can handle time series data, potentially with hidden nodes, and we benchmark them for the task of biochemical network reconstruction from time-resolved data on biochemical species’ concentrations against a rigorous random baseline. Specifically, in order to assess their performance against a known ground truth network, we apply them to synthetic data generated from models of biochemical kinetics of mass action and Michaelis-Menten type, simulated using stochastic differential equations (SDEs) from the chemical Langevin framework [36]. The stochastic trajectories generated in this way yield a particularly challenging setting for causal inference, due to strong autocorrelations, heterogeneous timescales, nonlinear interactions and noise terms with time, reaction rates and state dependencies. Our aim is to systematically evaluate how well causal discovery methods can reconstruct biochemical kinetic networks from complete or partially observed time series data, to identify the conditions under which they perform reliably, to highlight both their strengths and limitations, and thus to set the stage for their successful application to the study of intracellular biochemical reactions.

The article is organised as follows. We begin by benchmarking the three state-of-the-art causal discovery methods on synthetic time series data from two five-species toy models of biochemical kinetics, one with mass action and one with Michaelis-Menten dynamics. We then focus on tMIIC to investigate in depth how the relationship between sampling regimes and timescales of the system affects causal discovery performance. Next, we address the problem of partially observed networks, considering two scenarios: pure-noise latent confounders acting as common causes of observed species, and unobserved species that participate directly in the reactions of the network. Finally, we extend the benchmark to an experimentally calibrated model of the epidermal growth factor receptor (EGFR) signalling network, a well characterized biochemical system frequently subject to dysregulation in cancer, to demonstrate the feasibility of causal discovery on a realistic biochemical system.

### Box 1 Temporal causal discovery

#### Temporal causal networks

Temporal causal discovery constructs time-unfolded causal networks from time series data. These networks are called time series Directed Acyclic Graphs (ts-DAGs). Each variable is represented by a node at every time step, extending through the whole history of the dynamic system. This is the *full-time* network, and we call the set of nodes in a single time step a *layer* (Fig 1). Directed edges between nodes indicate a causal relation, which can connect two different variables (cross-link edges) or the same variable across time (auto-link edges). The number of layers between the causally related nodes denotes the *lag*, the time it takes for the cause to act. (Note that edges cannot go back in time as discussed below.) We say that a causal network constructed in this way is ‘time-unfolded’. By assuming that causal relations stay constant through time (the causal stationarity assumption), the full-time network may be reduced to the *window* causal network, a single unit of the full-time network that extends τ_max_ time steps into the past and is set to be invariant in time (Fig 1). τ_max_ is a hyperparameter supplied by the user and should be chosen with reference to the maximum expected lag in the system (for example by considering the decay of autocorrelations). Some algorithms (such as tMIIC [23]) allow a second hyperparameter Δ*t* to be chosen to determine the number of time steps between layers. The summary causal network is a ‘time-collapsed’ version of the network where an edge exists between nodes if there is an edge present at any time lag.

**Fig 1.**
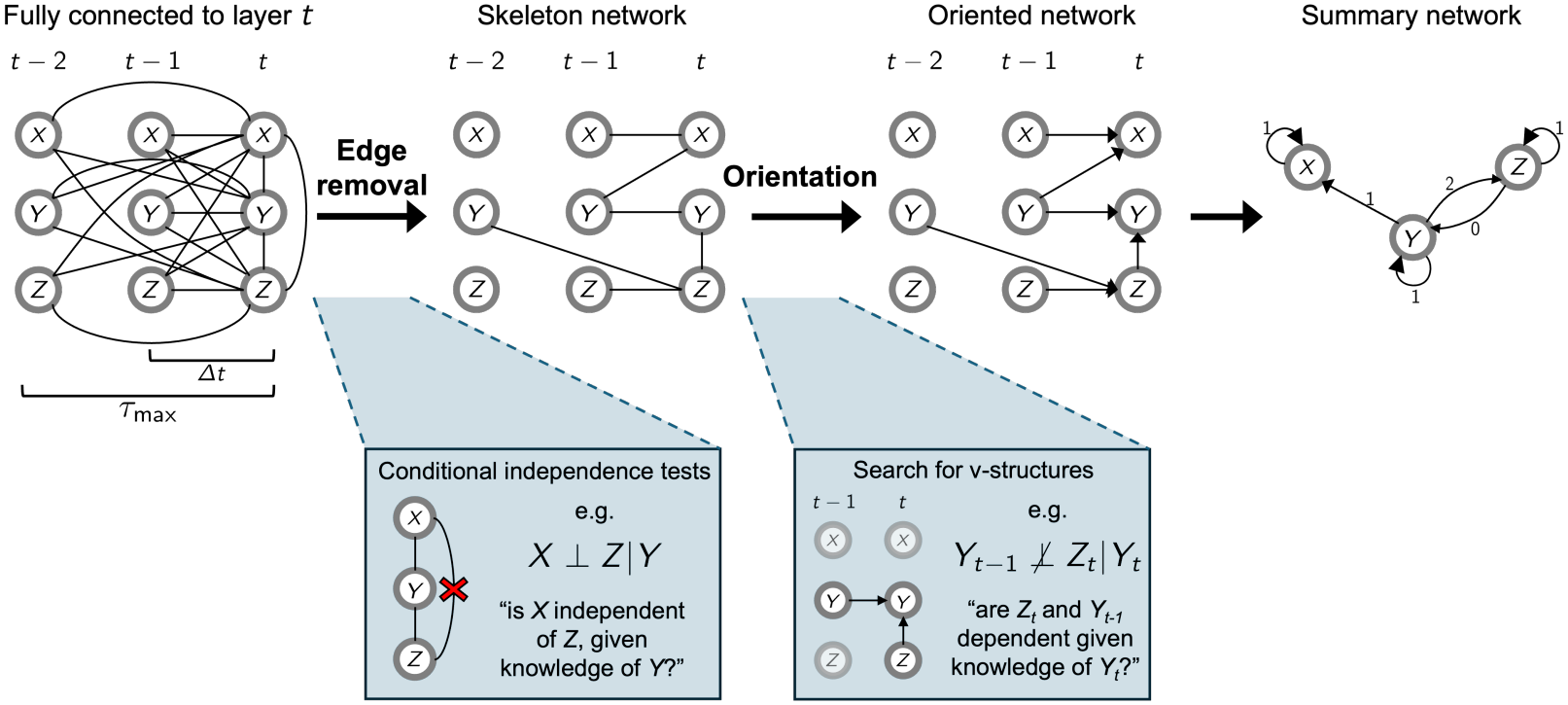
Overview of constraint-based causal discovery for temporal networks. Starting from a network fully connected to the contemporaneous layer, edges are removed with conditional independence tests (skeleton network) and then oriented, firstly by directing edges from the past into the future, and then using a v-structure search. The resulting time-unfolded network may also be represented as a compact summary network.

#### Constraint-based temporal causal discovery algorithms

Once a choice of τ_max_ and Δ*t* has been made, data may be input to a causal discovery algorithm to reconstruct the time-unfolded and corresponding summary causal network. Starting with a time-unfolded network fully connected to the present time layer (i.e., with every node in every layer connected to all nodes at time *t*), conditional independence between each node is tested, removing edges between nodes that are independent conditioned on some set of the other nodes. As in the simple example of Fig 1, the edge between a pair of nodes *X* and *Z* can be disposed of under the condition *X* ⊥ *Z* | *Y*, where ⊥ indicates independence given the conditioning variable *Y* . The exact mechanism for conditional independence testing depends on the algorithm used (see Methods for the algorithms under consideration here), but the aim is to produce a *skeleton* network for which edges between nodes imply a causal relationship.

The next phase is to orient the edges, giving them direction to show the flow of causality. For temporal causal networks, this is firstly done by directing any edges between layers into the future. Edges may be present between nodes at the contemporaneous layer (if dynamics occur more quickly than the time between the layers, or to indicate a potential hidden node). In this case, orientation rules from non-temporal causal networks are employed, whose key stage is to harness the *signature of causality* to identify *v-structures*. While in most cases the same conditional independence relations can correspond to multiple oriented structures (i.e., belong to the same Markov equivalence class), the relation 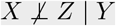 is a signature of causality because it exists in a class on its own and thus allows edges to be uniquely oriented as the v-structure *X* →*Y* ← *Z*. After orienting v-structures, edge directions may be *propagated* in a deductive manner under the constraints that: the network must be acyclic (causal networks are directed acyclic graphs); and v-structures exist if and only if the signature of causality has been observed. The final summary network is obtained by representing each variable by a single node for all time points and collapsing the edges in time. For an in-depth review of causal discovery for time series, see [22, 37].

## Benchmark study set-up

We built a rigorous benchmark for assessing the performance of causal discovery on time series data from stochastic biochemical systems, as visualized schematically in Fig 2. A model for a system of biochemical kinetics is defined with a set of chemical Langevin equations. From these governing equations, a ground-truth causal network is derived and time series data is generated via their numerical solution. This synthetic data (specifically, one stochastic trajectory per model) is then passed to the causal discovery methods, which infer a causal network compatible with the data. This predicted network is compared to the ground truth, and performance metrics are systematically computed. We use as main metrics of performance: precision, which measures how well the method is recovering the true network edges out of all its predicted edges; recall, which measures how well network edges are recovered out of all ground truth edges; and especially the F-score, which accounts for both precision and recall by computing their harmonic mean (see Methods for details). A high F-score (that is, close to 1) thus indicates that true edges are recovered while minimising false positives. We tested three state-of-the-art temporal causal discovery methods: tMIIC [23], PCMCI+ [20] (and its latent variable version LPCMCI [21]), and SVAR-FCI [24].

**Fig 2.**
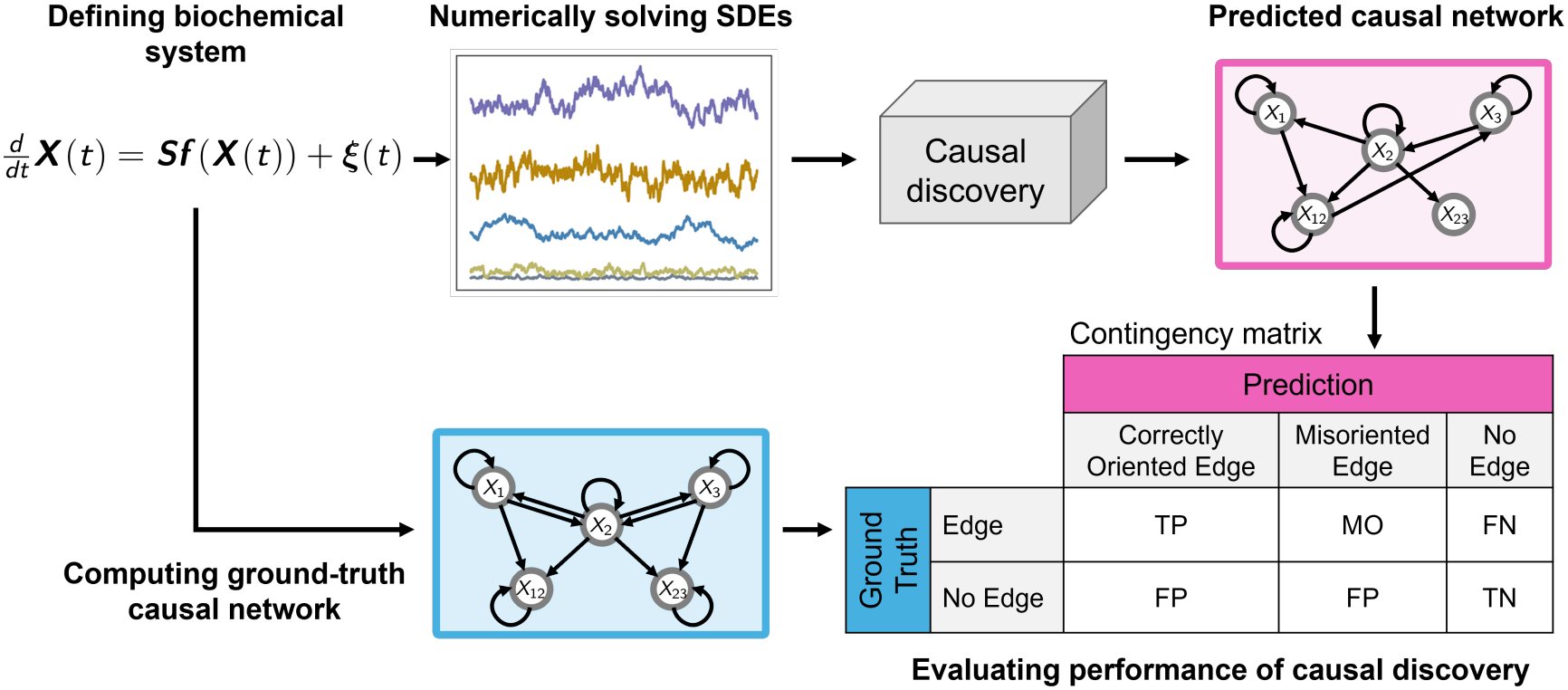
Overview of the benchmarking approach. The benchmarking approach of this study begins by choosing a model for biochemical kinetics in the form of a set of chemical Langevin equations. From these Stochastic Differential Equations (SDEs), the ground-truth causal network can be inferred. Data is generated by numerically solving the SDEs, to be passed to a temporal causal discovery method. The predicted causal network is compared to the ground truth to identify true positive edges (TP), true negatives (TN), false positives (FP), false negatives (FN), and misoriented edges (MO), and performance metrics are computed (see Methods).

### Chemical Langevin equations and corresponding causal networks

For each biochemical model we considered, the nonlinear and linearized systems were solved numerically to generate synthetic data. In the nonlinear case, we directly model the time evolution of *N* biochemical concentrations 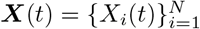 in a reaction volume *V* using Chemical Langevin Equations (CLEs) [38, 39]

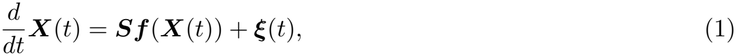

where ***S*** is the *N* × *R* (number of biochemical species × number reactions) stoichiometric matrix, and ***f*** is the *R*-dimensional flux vector containing the reaction rates. ***ξ*** is the *N* -dimensional vector of intrinsic noises which arises due to the stochasticity of reactions, to be interpreted according to the Itô convention [40]. It is Gaussian distributed with zero mean and covariance given by

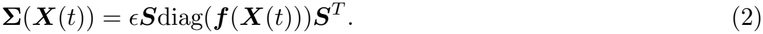

This noise is multiplicative due to the ***X***-dependent correlations (though with no temporal correlations). The parameter *ε* controls the scale of the noise variance and is the inverse of the reaction volume, *ε* = 1*/V* .

In the linearized case, we instead model fluctuations of the system’s concentrations *δ****X***(*t*) = ***X***(*t*) − ***Y*** around the steady state ***Y***, which is found by solving ***Sf*** (***Y***) = 0. The system of equations for these fluctuations is

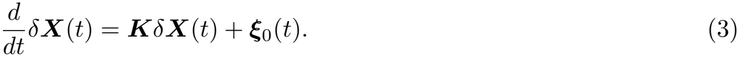

***K*** is the matrix of linearized reaction rates defined as

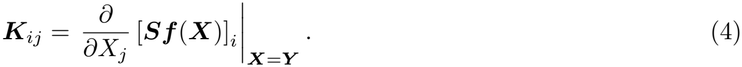

In these linearized dynamics, the noise ***ω***_0_(*t*) is additive with a constant-in-time covariance matrix,

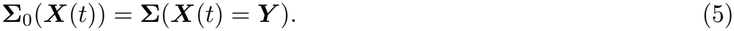

The adjacency matrix for the ground-truth summary causal network is constructed by setting non-zero elements of ***K*** to 1. To convert this to a time-unfolded network, we make the assumption that all edges have a lag of 1 (due to the Markovian nature of the dynamics), and that τ_max_ = 1 (see Box 1, Methods).

### Test cases: biochemical models with mass action and Michaelis-Menten kinetics

We considered two five-species toy models with prototypical biochemical reactions. The first model describes a system reacting under mass-action kinetics (Fig 3A), where species *X*_1_ and *X*_2_ form complex *X*_12_, and *X*_2_ and *X*_3_ form *X*_23_. These reactions are controlled by parameters representing the birth rates (*b*), reaction rates (*k*) and death rates (*d*).

**Fig 3.**
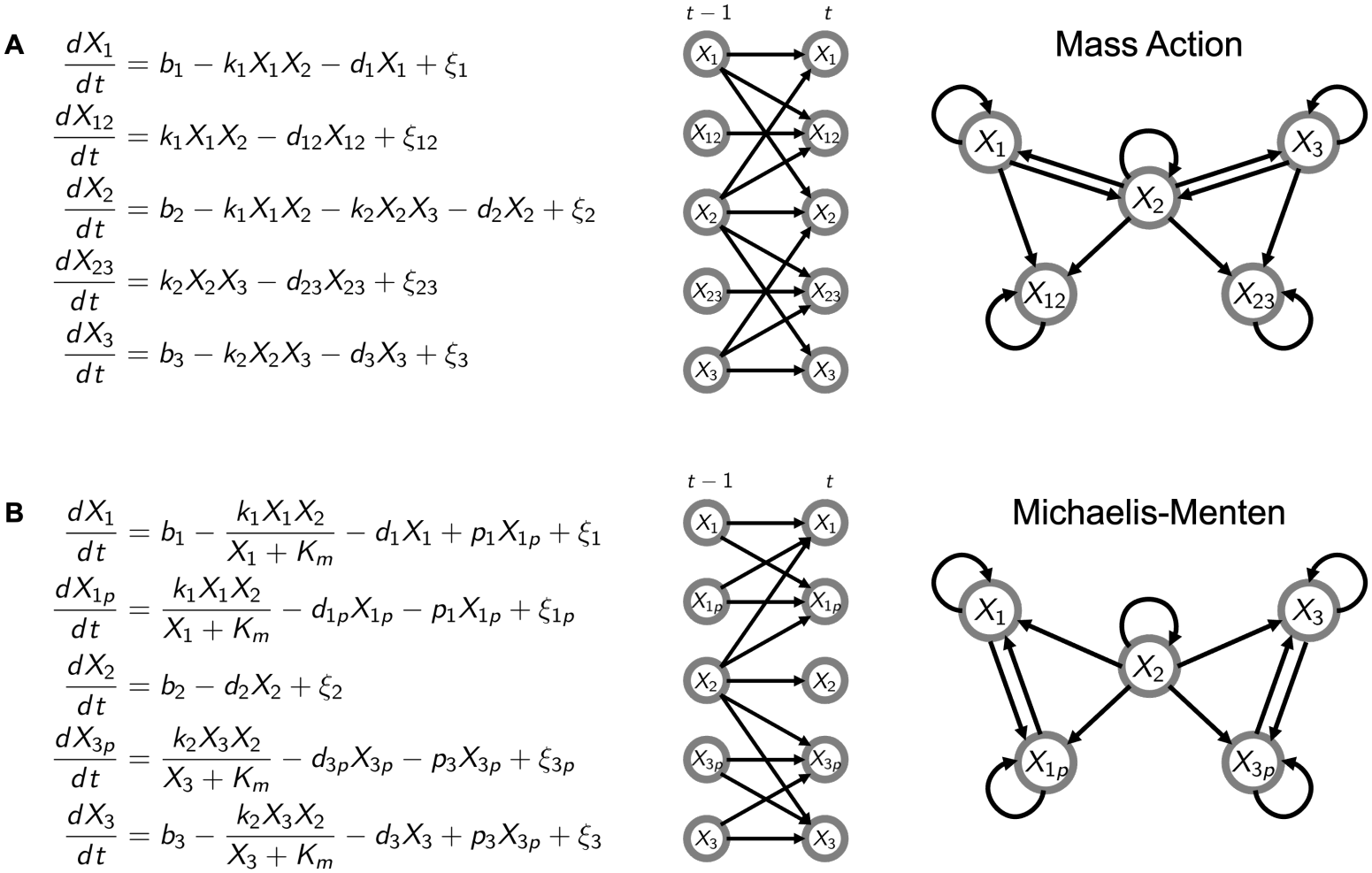
System of equations and corresponding causal networks for the five-species toy models. Chemical Langevin equations, where *ξ*_*i*_ is the multiplicative noise term associated with species *X*_*i*_ (left); time-unfolded causal network corresponding to the equations (centre); summary causal network corresponding to the equations (right). A: Five species model with mass action dynamics. B: Five species model with Michaelis-Menten dynamics.

The second model uses reversible Michaelis-Menten kinetics, with a similar but not identical network structure (Fig 3B). Here, proteins *X*_1_ and *X*_3_ are phosphorylated by an enzyme *X*_2_ to form *X*_1*p*_ and *X*_3*p*_, respectively. Enzyme-mediated phosphorylation adheres to a Michaelis-Menten functional form with Michaelis constant *K*_*m*_, while de-phosphorylation is controlled by parameters denoted by *p*.

For the two toy models, rate parameters were chosen by sampling from a log-uniform distribution, taking values between 0.1 and 10. 600 sets of parameters were generated in this way to cover a range of possible timescales for the system variables with diverse combinations of fast and slow, and homogeneous and heterogeneous dynamics.

To next test the methods on a more complex biochemical system, we considered the epidermal growth factor receptor (EGFR) signalling network. EGFR is in the family of protein-tyrosine kinase receptors and is involved in the control of cell growth, differentiation, and proliferation. For this network we used the model with experimentally calibrated parameters from [41], which has 23 variables and a mix of both mass action and Michaelis-Menten dynamics.

## Results

### Causal discovery performance on reconstructing biochemical networks from time-resolved data

Across all three causal discovery methods, performance improves, unsurprisingly, with the number of samples, (i.e., the trajectory length), which we denote by *M* (Fig 4A, top row). While edge counts increase with *M*, none of the methods approached the ground-truth edge count, even with 10^5^ samples (Fig 4A, second row), indicating that even with many samples all the three methods tend to underestimate the ground-truth sparsity. For PCMCI+ and SVAR-FCI, networks could only be inferred up to *M* = 1000 due to the high computational cost of the GPDCtorch kernel used for nonlinear data, which exceeded the 30-minute runtime limit we set per network prediction (Fig 4A, third row). The execution time for tMIIC is typically two orders of magnitude shorter than that of the other two methods. Consequently, the pairwise comparison in Fig 4B uses *M* = 1000 for all methods. For both the toy models and all methods auto-link edges are recovered with near perfect recall by *M* = 1000, whereas cross-link edges only begin to be detected when *M >* 100, and recall improves gradually after (Fig 4A, bottom row).

**Fig 4.**
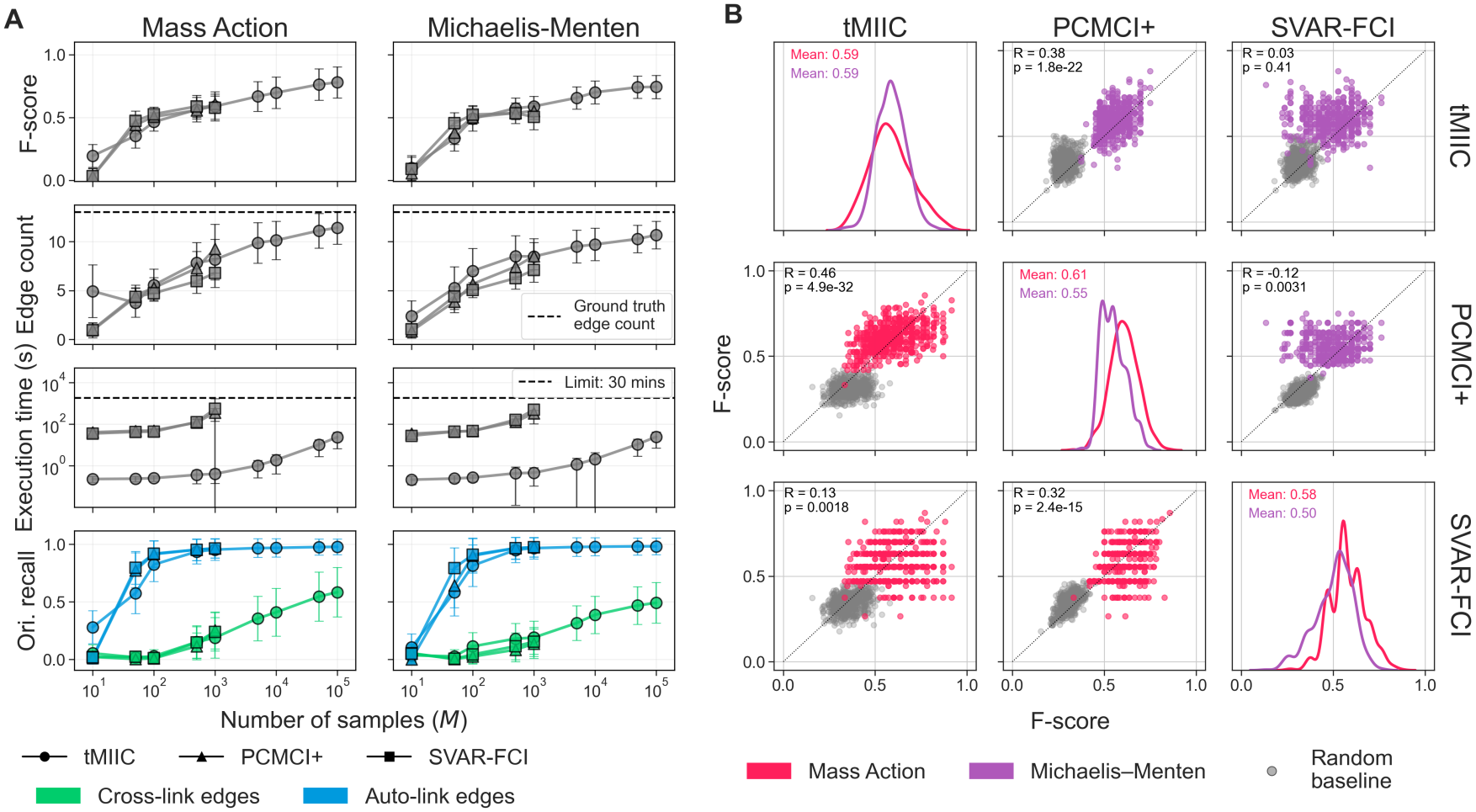
Performance of three causal discovery methods for biochemical network reconstruction for the five-species toy models. Nonlinear mass action and Michaelis-Menten models, Δ *τ* = 0.1. A) Mean F-score, edge count, execution time and orientation recall against number of samples *M*, across networks inferred from 600 parameter sets. B) Pairplot of F-scores for the three methods (*M* = 1000) across 600 parameter sets and both models. Diagonals show F-score density per method and model, off-diagonals show pairwise scatter with Pearson’s *R*.

For the mass action model, tMIIC performed best of the three with the very smallest sample size (*M* = 10), though this is not replicated to the Michaelis-Menten model. This likely reflects tMIIC’s ability to detect autocorrelation from limited data, as supported by the higher auto-link recall at this sample size.

Fig 4B presents a pair plot comparing F-scores across methods for *M* = 1000. The diagonal shows density distributions over parameter sets, highlighting variation in performance. SVAR-FCI exhibited the largest spread, with F-scores spanning almost the whole range from 0 (completely incorrect) to nearly 1 (fully correct). PCMCI+ is the method with the least cross-parameter set variation in performance (shown by the narrow peaks in the distributions) and the best mean F-score for mass action (0.61), but this is reduced to 0.55 for Michaelis-Menten. tMIIC has the same mean F-score of 0.59 for both models though a greater variability for mass action compared to Michaelis-Menten.

Off-diagonal scatter plots show per-parameter set correlations in method performance. For mass action, tMIIC and PCMCI+ and SVAR-FCI and PCMCI+ showed moderate, yet significant correlation (as measured by Pearson correlation coefficient R = 0.46 and R = 0.32, respectively, with *p <* 0.01 for both), while tMIIC and SVAR-FCI showed weaker association (R = 0.13, *p <* 0.01). For the Michaelis-Menten model, again there was an association between tMIIC and PCMCI+ (R = 0.38, *p <* 0.01), though comparisons against SVAR-FCI did not show significant correlations. Such comparisons highlight that the system’s reaction rates, and corresponding timescales, can constrain causal discovery predictive power regardless of the method.

To exhaustively quantitate the predictive power of the algorithms analyzed, here and in all the following numerical tests we compare it to a random baseline estimated following the methodology proposed in [42] for causal inference tasks and extending it to temporal causal discovery. For each network prediction, such random baseline consists of the mean performance of 100 randomly generated networks that preserve the sparsity of the prediction under consideration (see Methods). The scatter plots of Fig 4B show that, typically, the algorithms perform better than this random baseline, though there is more overlap of SVAR-FCI points with their random baseline than the other methods. Indeed on these toy models, SVAR-FCI on average shows the least improvement compared to the random baseline (S1 Table). Since SVAR-FCI performs only a little better than a random guess, it is perhaps to be expected that, when compared to methods which perform better, there is weak correlation performance-wise.

A similar pattern is observed for linearized dynamics, with network reconstruction generally improving with *M* (albeit SVAR-FCI’s mean F-score seems to plateau around 0.5 even with the largest *M*), and improved performance for PCMCI+ thanks to the feasibility of processing longer trajectories with the choice of kernel suitable for linear data (the ParCorr kernel, see S1 FigA). This observation also highlights a clear advantage of tMIIC ahead of the other two methods tested, that is, its agnosticism to the type of the data-generating dynamics. PCMCI+ and SVAR-FCI require a choice of kernel for independence tests according to whether the dynamics is linear or nonlinear, whilst this is not the case for tMIIC.

### Causal discovery is sensitive to data sampling regimes relative to dynamical timescales

Based on the comparisons of three causal discovery methods made, we chose tMIIC as the method for which to continue further investigations into the impact of the data-generation and sampling processes on performance. We have seen that, out of the methods tested, it was able to infer causal networks from linearized as well as nonlinear data, handling many samples efficiently and with reasonable recall and precision.

The density plots in Fig 5A show the distribution of F-scores for network reconstructions for the mass action models for data sampled at three different sampling intervals (Δ _*τ*_) for fixed trajectory length *M* . It appears that sampling the data less frequently (Δ _*τ*_ = 10^−1^) improves the average performance across the 600 parameter sets, but there is some overlap: some networks sampled at Δ _*τ*_ = 10^−3^ outperform other sampled at Δ _*τ*_= 10^−1^.

**Fig 5.**
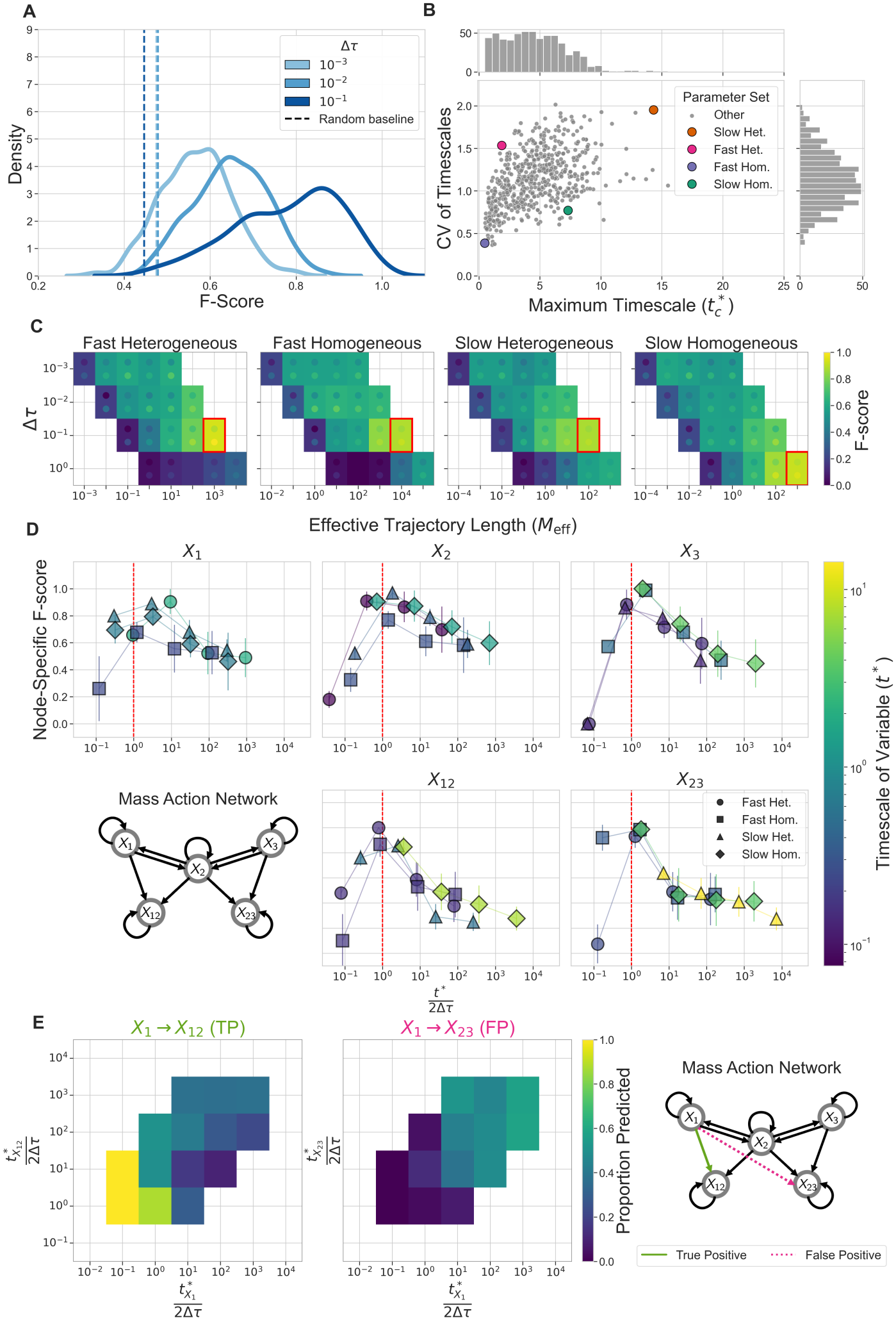
Performance of network reconstruction by tMIIC with changes to the sampling interval of the data. Nonlinear mass action models, *M* = 100, 000. A) F-score density across 600 parameter sets at three values of Δ _*τ*_. B) Characteristic timescale (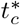, maximum relaxation time across the five timescales) against timescale heterogeneity (coefficient of variation, CV) for each parameter set, with four selected parameter regimes (fast/slow, homogeneous/heterogeneous) highlighted. C) Mean F-score heatmaps for the four selected regimes, varying *M*_eff_ and Δ _*τ*_, averaged over 10 realizations. Dots mark ± 1 SD. D) Node-specific F-scores against *t*^*\**^*/*2Δ _*τ*_ for each variable. Red dashed line shows sampling interval equal to Nyquist rate (*t*^*\**^*/*2Δ _*τ*_ = 1). E) Proportion of predicted edges as a function of *t*^*\**^*/*2Δ _*τ*_ for both variables in a representative true positive edge (left), and representative false positive edge (right), across all 600 parameter sets and three values Δ _*τ*_ = 10^−3^, 10^−2^, 10^−1^ (1800 networks total). Bins with *<* 10 observations omitted (4 excluded for the true positive edge, 2 for the false positive edge).

To investigate this observation further, we examined how the timescales of the system interact with the sampling interval (Δ _*τ*_). We first defined a system’s characteristic timescale 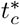 as the slowest (i.e., the maximum) relaxation timescale across the species in the system (see Methods for details). Next we introduced a quantity, *M*_eff_, which we refer to as ‘effective trajectory length’, to quantify the number of effectively independent relaxation time steps in a trajectory with *M* samples and sampling interval Δ _*τ*_ given the system’s characteristic timescale 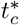

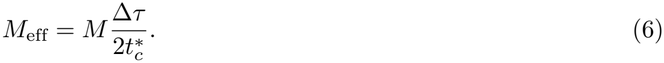

The definition of *M*_eff_ accounts for the fact that successive samples are correlated over the characteristic timescale 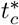. If 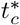 is large compared to Δ _*τ*_, many data points carry redundant information and the effective trajectory length is reduced [43], because the whole trajectory *M* Δ _*τ*_ samples adequately only a few relaxation times. However, if the time between samples (Δ _*τ*_) increases, the samples carry more independent information, and the effective trajectory length increases.

Four parameter sets were selected for each combination of fast/slow characteristic timescale and homogeneous/heterogeneous timescales across variables, see Fig 5B. Fig 5C presents the F-scores of networks inferred from simulations of these four parameter sets, varying Δ _*τ*_ and the effective trajectory length *M*_eff_. In line with results shown in Fig 4A, performance increases with *M*_eff_ for all parameter sets and all values of Δ _*τ*_, confirming, as expected, that sampling data over many relaxation times is more informative. For both fast systems, the optimal Δ _*τ*_ is 10^−1^, while the slow homogeneous system performs best at Δ _*τ*_ = 10^0^. The slow heterogeneous example favours 10^−1^, which is likely due to the fact that only one variable has a slow timescale (on the order of 10), while the others are faster (on the order of 0.1). However, the slow homogeneous example has four variables with a timescale on the order of 1 and one on the order of 0.1.

These results indicate that global network reconstruction performance is most accurate when the sampling interval is consistent with the timescales of the system.

Node-specific performance also requires sampling intervals of the data to be consistent with timescales of the variables. The Nyquist-Shannon sampling theorem [44] states that a signal can be faithfully reconstructed if it is sampled at least twice as fast as the signal’s component with the highest frequency. Recast in terms of times, the Nyquist theorem implies optimal signal reconstruction when the sampling interval Δ _*τ*_ approaches half of its typical relaxation timescale, *t*^*\**^*/*2 in this setting. In Fig 5D, we indeed observe that the node-specific F-score peaks when the sampling rate approaches the Nyquist rate, that is, when *t*^*\**^*/*2Δ _*τ*_ is close to 1. Notably, performance decreases more sharply for *t*^*\**^*/*2Δ _*τ*_ *<* 1 (below the Nyquist rate) than for *t*^*\**^*/*2Δ _*τ*_ *>* 1 (above the Nyquist rate), showing that oversampling is less detrimental than undersampling.

Because each edge involves two nodes, we also investigated the interaction between the timescales of the biochemical reaction coupling two nodes, and the frequency of prediction of true positive and false positive edges. In Fig 5E we plot heatmaps showing the proportion of predicted edges (across all 600 parameter sets and Δ _*τ*_ = 10^−3^, 10^−2^, 10^−1^) with combinations of *t*^*\**^*/*2Δ _*τ*_ for both nodes in the edge. True positive edges are more frequently predicted in the region where *t*^*\**^*/*2Δ _*τ*_ is close to 1 for both variables whereas false positives are more common when *t*^*\**^*/*2Δ _*τ*_ is much larger than 1.

### Causal discovery can detect the presence of hidden variables

#### Causal networks with hidden nodes

We next considered the case where the dynamics are partially observed. To model this scenario, we partition the system into a *subnetwork* of observed nodes and a *bulk* of unobserved, hidden variables. The subnetwork-reduced SDEs describing the time evolution of the subnetwork under the influence of the unobserved bulk species can be obtained by coarse-graining (i.e., marginalizing) over the bulk time evolution and are tractable exactly in the linearized dynamics case (see [10–12], Methods). This is hence the case to which we stick for assessing causal discovery’s ability to detect hidden nodes and their typical timescales.

The linearized rate matrix ***K*** may be as well divided into four blocks

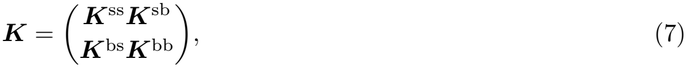

where ***K***^ss^ and ***K***^bb^ represent biochemical interactions purely within the subnetwork and bulk, respectively, while ***K***^sb^ and ***K***^bs^ are the cross-interactions between the observed and hidden variables.

The key observation in this regard is that only the subnetwork nodes which have a hidden node as a cause will receive additional edges, and, following [10–12], will be referred to as *boundary nodes*. These correspond to any variable with a non-zero entry in its row of ***K***^sb^.

Boundary nodes receive additional edges of two distinct types: one if the hidden node is a confounder (resulting in contemporaneous edges which may be bidirected if the hidden node has no autocorrelations), and the other if the hidden node is a direct intermediate between two observed subnetwork nodes. In the latter case, the matrix multiplication ***K***^sb^***K***^bs^ yields an *N* ^s^× *N* ^s^ adjacency matrix for the additional subnetwork edges with lag *>* 1 (i.e., its non-zero elements indicate delayed edges between subnetwork variables).

Characteristic timescales (and amplitudes) may be associated to causal delayed edges, since in the context of subnetwork-reduced dynamics obtained by coarse-graining over the hidden nodes, delayed effects can be modelled through non-Markovian terms, i.e., terms that connect the current subnetwork state to previous time steps (see [10–12] and Eq 14 in Methods). Specifically, the first of these terms is a memory function, which for the linearized equations is given by:

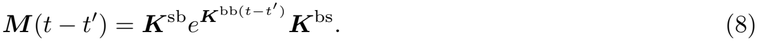

This function explicitly accounts for the influence of the subnetwork on the bulk at time *t*^*’*^ that is subsequently fed back into the subnetwork at a later time *t* (see Methods). The second non-Markovian term is an effective noise term that is ‘coloured’, i.e., it displays correlations over time (see Eqs 14, 15, and 16 in Methods).

Generally, it is the slow reaction timescales in the unobserved bulk that lead to long decay times of these bulk-induced, non-Markovian effects in the subnetwork. This can be explicitly appreciated through an estimate of timescale of the memory function’s decay that can be calculated as proposed in [10] (see Methods). For the self-memory terms (i.e., terms capturing non-Markovian effects of a subnetwork variable *s* into itself at later times) the explicit expression for the memory timescale reads

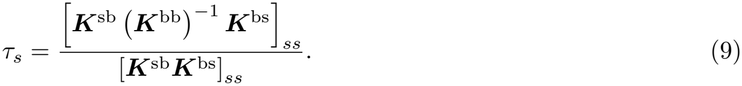

that is, *τ*_*s*_ is a weighted average of elements of (***K***^bb^)^*−1*^, and thus scales with the inverse of the reaction rates within the bulk (contained in ***K***^bb^).

#### Benchmark of causal discovery of hidden nodes

In Fig 6 we present results from a partially observed network of the Michaelis-Menten model; we report analogous results for mass action in S3 Fig. To investigate the effect of both types of hidden node (one inducing delayed edges, the other contemporaneous edges), we treated one of the original five variables in the toy models (*X*_3_) as the unobserved bulk and we augmented them with an additional time-uncorrelated confounder *η*. This is a variable modelled by pure white noise acting as a common cause of *X*_1*p*_ and X_3*p*_ in the case of the Michaelis-Menten model (Fig 6A) and of *X*_12_ and *X*_23_ for the mass action one (S3 FigA).

**Fig 6.**
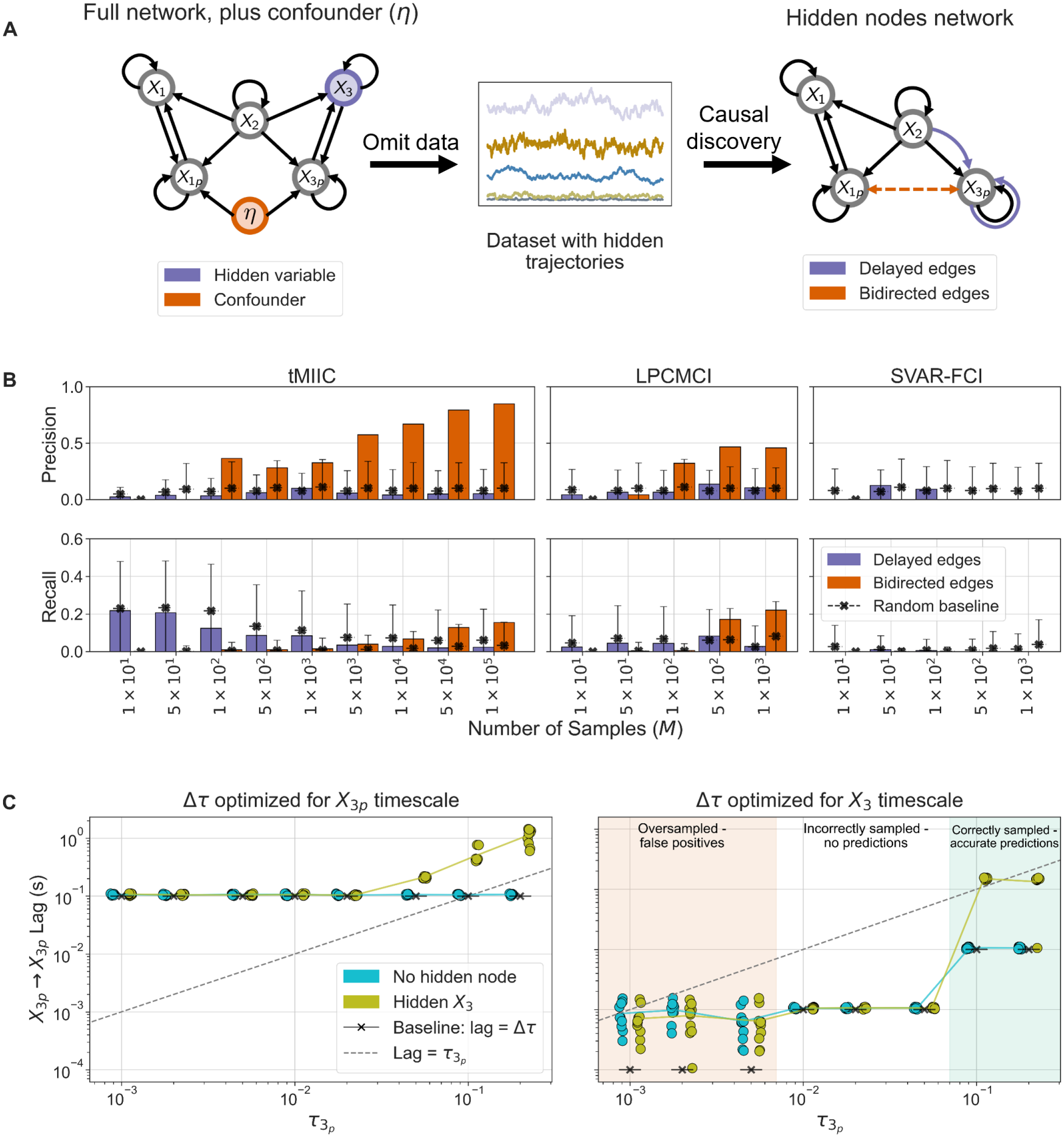
Summary of additional edges predicted with hidden nodes. A) Full network with uncorrelated confounder *η* (orange), and hidden variable *X*_3_ (purple). Hidden trajectories are withheld from causal discovery (centre), to obtain the reduced causal network (right). The hidden node induces delayed edges (purple) and bidirected edges (orange) are induced by the confounder. B) Precision and recall of the delayed and bidirected edges indicating hidden nodes across sample sizes *M*, for tMIIC, LPCMCI and SVAR-FCI (nonlinear data, Δ*τ* = 0.1, *τ*_max_ set according to the slowest timescale in the subnetwork, GPDCtorch kernel for LPCMCI and SVAR-FCI, maximum *M* limited by 30-minute limit for computation). Black crosses mark random baseline values. C) Lag of the predicted *X*_3*p*_ → *X*_3*p*_ edge against hidden node timescale (*τ*_*s*_), for one parameter set (recall = 1.00, precision = 0.93), with *d*_3_ varied to control *τ*_*s*_, 10 random realizations, linearized model (points jittered to enhance visualization). Dashed line represents lag = *τ*_*s*_, and crosses are the baseline Markovian (lag 1) edge. Left: Δ*τ* set according to the boundary species (*X*_3*p*_) timescale. Right: Δ*τ* set according to the hidden node (*X*_3_) timescale.

When the confounder *η* and the trajectory for the variable *X*_3_ are held back from the causal discovery algorithms, we expect to infer a network for the reduced system of *X*_1_, *X*_1*p*_, *X*_2_ and *X*_3*p*_ (the subnetwork) with additional contemporaneous bidirected edges and delayed edges (Fig 6A, right), carrying the signature of hidden nodes. The ground truth, subnetwork-reduced causal network can be derived by identifying inducing paths between its nodes with respect to the hidden nodes, which result in one bidirected edge and two delayed edges (see Methods for more details).

Fig 6B measures the quality by which the additional edges are predicted through precision and recall of the specific bidirected and delayed edges we expect in this network. For tMIIC at a low number of samples, recall of delayed edges is around 0.2, close to the random baseline. Bidirected edges are not predicted until *M* = 500, and after this point the frequency increases as the number of samples increases. Precision is high, suggesting that when the bidirected edge is predicted there are few false positives, which makes the confounder easy to place. For LPCMCI (the latent variable version of PCMCI+ [21]), we observe a similar pattern for bidirected edges, and by *M* = 1000 (the maximum number of samples that could be processed by this method in the 30-minute time limit) the bidirected edge recall exceeds that of tMIIC at *M* = 100, 000 samples. Recall and precision for delayed edges are typically poor (close to the random baseline) for LPCMCI. SVAR-FCI does not perform well at this task, never predicting bidirected edges, or reliably predicting delayed edges. Since SVAR-FCI uses stationarity to remove edges whose separating set extends beyond *τ*_max_, edges which are fully contained within the time window may be incorrectly removed. This in turn affects further independence tests, and bidirected edges are not inferred.

We performed the same analysis on the mass action model, augmented with a confounder *η* acting on the protein complexes *X*_12_ and *X*_23_. The resulting subnetwork when *X*_3_ and *η* are hidden is shown in S3 FigA. Recall and particularly precision of both types of edge are above the random baseline for tMIIC and LPCMCI with sufficient *M* (S3 FigB). For SVAR-FCI, both of these quantities are close to zero for the bidirected edge for all tested values of *M*, while recall for the delayed edges remains low.

Based on these performance results, we restricted to tMIIC to investigate the relationship between the lag of the hidden node-induced edges in the causal network and the timescale of the non-Markovian effects in the subnetwork-reduced SDEs. Since one has an explicit expression for the self-memory timescale for the linear dynamics (Eq 9), we focus on the auto-link delayed edge *X*_3*p*_ → *X*_3*p*_ in the linearized model. When calculating the timescale of the self-memory on the subnetwork species *X*_3*p*_ for the model with hidden nodes of Fig 6A, Eq 9 reduces simply to *τ*_3*p*_ = 1*/K*^bb^ and with all other parameters fixed scales with 1*/d*_3_ (see Methods), that is, the memory’s timescale is fully controlled by the decay rate *d*_3_ of the bulk species *X*_3_. Similarly, the correlations of the effective coloured noise on *X*_3*p*_ have a decay time set by 1*/K*^bb^, thus *τ*_3*p*_ recapitulates all the timescales associated to non-Markovian effects. To test whether causal discovery can capture such timescale through its delayed edges, we selected a single parameter set with a high memory amplitude and altered the decay rate *d*_3_ to enforce *τ*_3*p*_ on a scale from 1 × 10^−3^ to 2 × 10^−1^. For each value of *τ*_3*p*_ we generated 10 sets of trajectories with different random seeds and passed them to tMIIC with *τ*_max_ = 15. In these tests, the choice of Δ*τ* is particularly important. Firstly, due to the importance of choosing Δ*τ* with respect to the timescales of the system to optimize performance (Fig 5). Secondly, now that we are searching for edges with specific lags to detect hidden nodes, it is also important to ensure that we sample the data in a way that allows the window causal network (see Box 1) to cover the region over which we expect the delayed edge to occur. Of course, in practice, if data is hidden, the timescales of these unseen components are typically unknown. A sensible choice (and the one that appeals to the first consideration) is to sample the data according to the timescale of the boundary nodes, which in this case is just *X*_3*p*_. In Fig 6C (left), we show the results of these tests with Δ*τ* = 0.1, the optimal choice for the timescale of *X*_3*p*_. When there is no hidden node, the baseline expectation is to infer only a Markovian edge *X*_3*p*_ → *X*_3*p*_, i.e., an edge with lag = Δ*τ* – indeed this is the case for all values of *τ*_3*p*_. However, when the *X*_3_ node is hidden, we do see the prediction of delayed edges from *X*_3*p*_ to itself when *τ*_3*p*_ is 0.05 or greater. At the smaller values of *τ*_3*p*_, where the dynamics of the hidden node are faster, the resolution of the time unfolded network (with steps of Δ*τ* = 0.1 between each layer) is too coarse for tMIIC to detect the effect of the hidden node on *X*_3*p*_ separately from the Markovian effect of *X*_3*p*_ on itself.

We carried out a second version of these tests, choosing Δ*τ* in such a way as to maximize the ability to detect delayed edges with Δ*τ* 10 times smaller than *τ*_3*p*_, so that *τ*_3*p*_ lies within the range up to *τ*_max_. The result of this is three choices of Δ*τ* across the range of chosen *τ*_3*p*_, as shown in Fig 6C (right), each displaying different behaviours highlighted with different shaded regions in the plots. For the smallest values of *τ*_3*p*_, on the order of 10^−3^, we chose Δ*τ* = 10^−4^. In this regime, delayed edges are predicted for networks both with and without hidden nodes. This corresponds to oversampling: Δ*τ* is much faster than the timescales in the system and we observe false positive edges, aligning with the results shown in Fig 5E. For the values of *τ*_3*p*_ on the order of 10^−2^, we chose Δ*τ* = 10^−3^. In this region, we now do not see the false positive delayed edges when there is no hidden node, but also do not see them when *X*_3_ is hidden. This Δ*τ* is still not compatible with the timescales of the subnetwork, so while we do not observe the same false positive edges as with the smaller Δ*τ*, the networks are in general not well reconstructed and the specific edge we are looking for cannot be detected, perhaps due to other inaccuracies within the network. For *τ*_3*p*_ on the order of 10^−1^, Δ*τ* = 10^−2^. For these networks, we do not observe a false positive delayed edge when there is no hidden node, but now do when *X*_3_ is hidden. This region is well predicted due to the choice of Δ*τ* which is more compatible with the timescales of the subnetwork than the prior choices, allowing for a more accurate overall network prediction. With these results we show that with a careful choice of Δ*τ* and *τ*_max_ that accounts for both the subnetwork and bulk, tMIIC can infer the presence and timescale of the hidden node.

### Application to the epidermal growth factor receptor signalling network

We next applied these causal discovery methods to the model for the Epidermal Growth Factor Receptor (EGFR) signalling pathway from [41]. This model contains 23 species and 25 reactions of mass action and Michaelis-Menten type, leading to a causal network with 23 nodes and 274 edges (visualized in Fig 7A). The variables in this model are highly heterogeneous in terms of node degree and of timescales which span three orders of magnitude (Fig 7B). The model was passed through the computational pipeline described in Fig 2, using the experimentally calibrated parameter values of the original paper.

**Fig 7.**
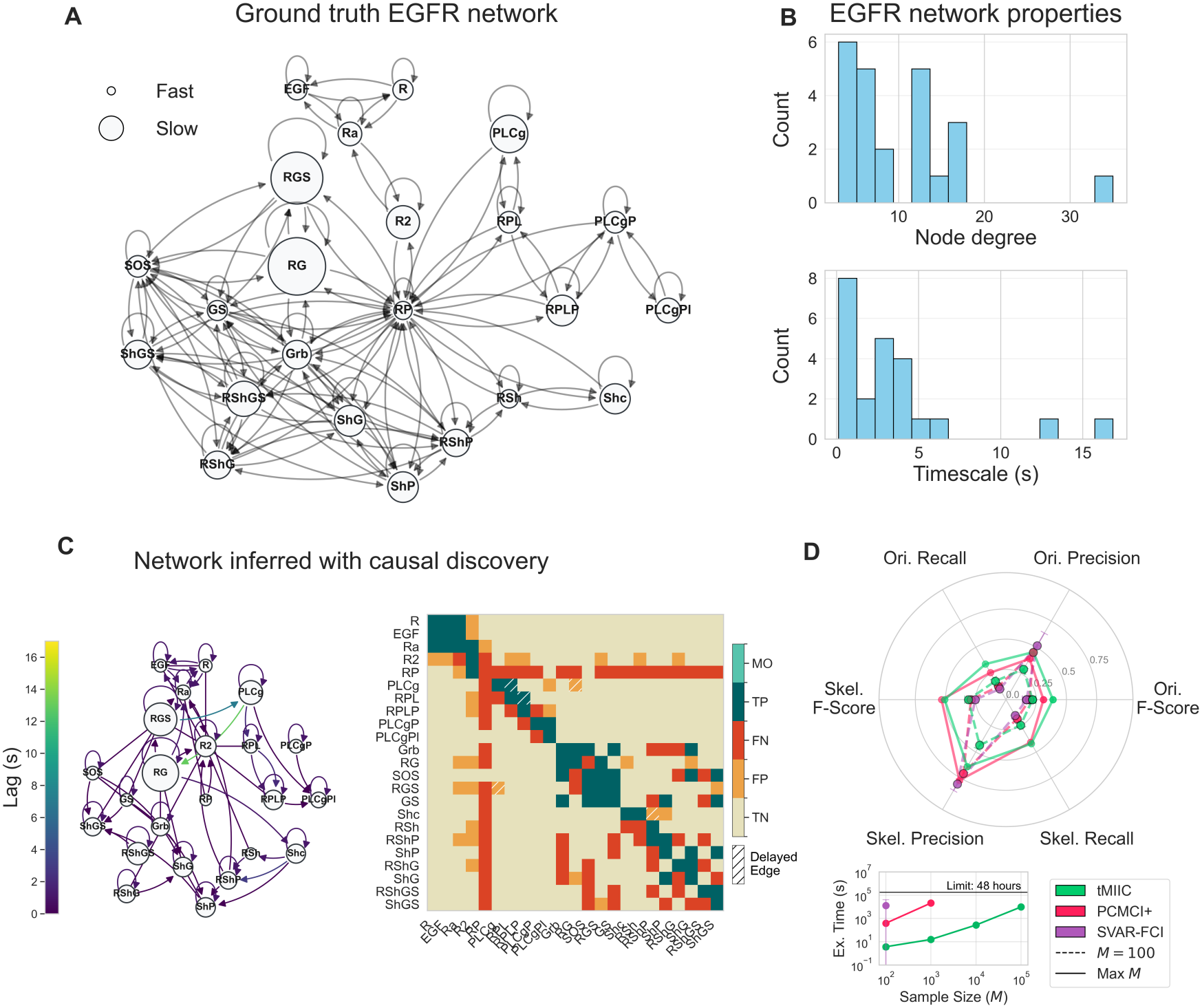
Results of causal discovery on the EGFR network. A) Summary causal network for the EGFR model, derived from the kinetic equations. B) Histograms of node degree and timescales across the network’s species. C) Example network and time-collapsed heatmap representation learnt by tMIIC. Nonlinear model, Δ*τ* = 1, *τ*_max_ = 17, *M* = 100, 000. D) Radar plot comparing the three methods’ performance. Dashed lines show performance metrics at *M* = 100, the maximum common sample size, while solid lines show performance at maximum feasible *M* (1000 for PCMCI+ and 100,000 for tMIIC). Inset shows execution time against sample size. Nonlinear data, Δ*τ* = 1, *τ*_max_ = 17, GPDCtorch kernel for PCMCI+ and SVAR-FCI.

We present an example network predicted by tMIIC in Fig 7C, choosing Δ*τ* = 1 to match most of the timescales of the system (see S4 Fig) and with *τ*_max_ = 17 so that the slowest timescale in the system can be captured. The heatmap is a time-collapsed representation of the network and shows that a reasonable level of prediction quality is achieved. Certain modules of the network are particularly well predicted (one involving EGFR, R, Ra, and another involving Grb, RG, SOS, RGS and GS), and most non-edges are true negatives, i.e., correctly not predicted. However, the species RP (the phosphorylated EGFR), is involved in many false negative edges. RP has the fastest timescale in the system (on the order of 0.1) and since we have chosen a sampling regime in accordance to the average timescales of the system (on the order of 1, see S4 Fig), RP is left undersampled. As shown in experiments on the toy models, this drastically reduces node-specific performance (Fig 5D). RP is also the node with the highest degree in the ground truth network (degree = 35, Fig 7B).

We next compare tMIIC to PCMCI+ and SVAR-FCI and plot time-collapsed performance metrics along with execution times for various sample sizes (*M*) in Fig 7D. With a time limit of 48 hours, SVAR-FCI (using the GPDCtorch kernel) could handle up to *M* = 100, PCMCI+ *M* = 1000 (also using the GPDCtorch kernel) and tMIIC *M* = 100, 000 samples (Fig 7D, inset). SVAR-FCI achieves high precision, both at the network skeleton (skel.) and edge orientation (ori.) level prediction, though this comes at the cost of poor recall, indicating that overall few edges are predicted and resulting in the lowest F-scores of the methods compared. As expected, with their higher maximum sample sizes tMIIC and PCMCI+ performance is improved. PCMCI+ outperforms tMIIC in terms of the skeleton metrics, though when considering orientation, tMIIC is marginally better.

We compare two subnetworks of this model: the first of these (Fig 8A) is the subnetwork studied in [10–12], motivated by placing the module involving the protein Shc and all complexes it is involved in into the bulk, and which we refer to as the *arbitrary subnetwork*. It comprises 15 nodes, four of which are boundary species (SOS, RP, Grb and GS). The second (Fig 8B) is instead chosen according to the timescales, taking the fastest species into the subnetwork and leaving the slowest in the bulk. Defining ‘fast’ as *t*^*\**^ *<* 1.5*s* gives a *fast subnetwork* of nine nodes, seven of which are boundary. This split is motivated by our earlier finding that performance strongly depends on the relationship between Δ*τ* and the timescales (Fig 5).

**Fig 8.**
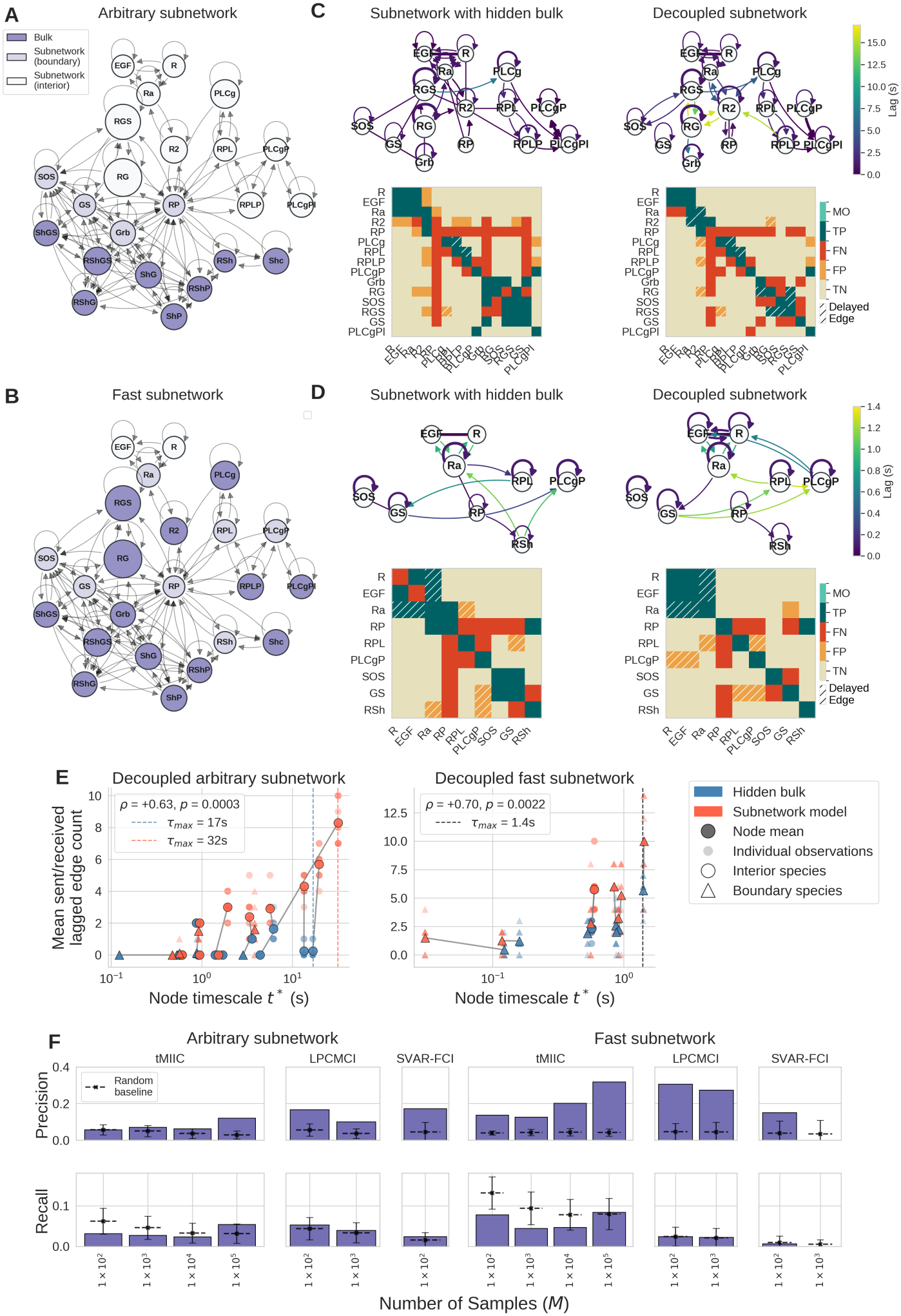
Results of causal discovery on the EGFR network with hidden nodes. A) Arbitrary subnetwork, with Shc and its complexes as the bulk (dark purple), and four boundary nodes (light purple). B) Fast subnetwork, with the slowest species as the bulk (dark purple), and seven boundary nodes (light purple). C) Representative tMIIC predictions for the arbitrary subnetwork (top), with adjacency matrix heatmaps comparing to the ground truth (bottom). Nonlinear model, Δ*τ* = 1, *τ*_max_ = 17, *M* = 100, 000. D) As C, for the fast subnetwork. Nonlinear model, Δ*τ* = 0.1, *τ*_max_ = 1.4, *M* = 100, 000. E) Variable timescale against mean number of lagged edges per node, with Spearman’s *ρ* and p-value *p* assessed using a permutation test with edge count values shuffled over 10,000 permutations. F) Precision and recall of lagged edges (indicating the hidden nodes) for the three methods on arbitrary (left) and fast (right) subnetworks. Black crosses show mean random baseline values. Error bars indicate ±1 SD.

As in the toy models, we omit trajectories for data that is to be ‘hidden’. We chose Δ*τ* and *τ*_max_ according to the timescales of the corresponding subnetwork as for the general user of causal discovery, this would be the only information available. For the arbitrary subnetwork this was Δ*τ* = 1.0 and *τ*_max_ = 17, and for the fast subnetwork Δ*τ* = 0.1 and *τ*_max_ = 1.4. We also constructed a system to act as a control, where the bulk is removed from the dynamics entirely. In these *decoupled* subnetworks, we clamp the bulk variables to their long-time steady-state values, making the subnetwork effectively dynamically independent of the bulk. In this case, there are no hidden nodes, hence no non-Markovian dynamics due to memory effects, so our hypothesis is that neither should yield lagged edges.

From the ground-truth network with hidden nodes, we expect causal discovery to predict lagged edges into each of the boundary nodes. From our tests on the arbitrary subnetwork we observe that lagged edges are not predicted into these nodes, and few lagged edges are predicted overall (Fig 8C, left). In our tests of five-species toy model (Fig 6C), we chose a specific parameter set with a high memory amplitude to maximise the chance of predicting the lagged edge. In this model, we use experimentally calibrated parameters, and for the arbitrary subnetwork, the strongest memory amplitude is on the order of 0.1 which is likely insufficient to allow detection by tMIIC (S6 FigA, left). Additionally, the timescales in the arbitrary subnetwork remain heterogeneous, so a single Δ*τ* cannot be near the optimal Nyquist-rate-related sampling interval for all species simultaneously. These two factors which were identified as crucial in the toy models therefore fail, so this absence of edges indicating the hidden nodes is perhaps not surprising.

The fast subnetwork was chosen to improve on these two aspects of insufficient memory amplitudes and heterogeneous timescales. The heatmap of amplitudes (S6 FigA, right) shows that some memories now have amplitudes on the order of 1.0. In particular, the amplitude relating to the *RP → RP* is 7.92, and some amplitudes involving *RSh* are also substantial. The predictions show a slight improvement compared to the arbitrary subnetwork (Fig 8D, left). As in the fully observed examples (Fig 7C), we again see systematic false negatives with edges involving *RP* which is most likely due to the fact that it remains the fastest species in the subnetwork and therefore the most undersampled for the chosen Δ *τ* . A large memory amplitude alone is thus not sufficient if the receiving node is sampled too far from its Nyquist rate. Lagged edges do appear in association with *RSh*, a node which has some strong memory amplitudes and timescales closer to Δ*τ* . These are not the specific edges from the ground truth, so we cannot count them as true positive, but their location suggests that tMIIC may be detecting some signal of the slower bulk through this node.

Despite our hypothesis that the decoupled subnetworks should not contain lagged edges since there are no hidden components to induce non-Markovian effects, we in fact observe that in these subnetworks there are more lagged edges predicted than in the subnetworks with hidden nodes (Fig 8C, right and Fig 8D, right). To investigate this, we compared timescales in the decoupled models to those in the full dynamic model (S6 FigB), observing that in some cases timescales are altered by an order of magnitude. Removing the bulk does not necessarily leave the subnetwork dynamics unchanged, but can in fact induce (in some cases) extreme effects on timescales of the remaining subnetwork species. These changes leading to even longer timescales could explain why lagged edges are predicted in these decoupled subnetworks.

In Fig 8E, we show the positive correlation between a node’s timescale and the number of predicted lagged edges it is associated with (Spearman *ρ* = 0.63, *p* = 0.0006 for the decoupled arbitrary subnetwork; *ρ* = 0.70, *p* = 0.0014 for the decoupled fast subnetwork). Nodes that are slower relative to Δ*τ* accumulate lagged edges regardless of whether or not there are hidden nodes. In this system, the result we characterised in the toy models (that lagged edges are associated with hidden nodes, and these lags correspond to timescales of the hidden effects) is now confounded by the more complex properties of this real-world network.

We also make comparisons between tMIIC, LPCMCI, and SVAR-FCI, scoring precision and recall of the prediction of edges associated with the hidden nodes in the time series Maximal Ancestral Graph (ts-MAG). In the arbitrary subnetwork (Fig 8F, left), precision remains low for all three methods and sample sizes, though typically at or exceeding the random baseline. At the smaller sample sizes, LPCMCI and SVAR-FCI outperform tMIIC on precision. Recall is also low across methods (below 0.1) and sample sizes, though tMIIC with 100,000 samples does exceed its random baseline performance. For the fast subnetwork (Fig 8F, right), performance improves for tMIIC, precision increasing with sample size from around 0.1 at *M* = 100, to around 0.35 at *M* = 100, 000, far exceeding the random baseline for all sample sizes. Precision for LPCMCI is also far above the baseline, though SVAR-FCI loses all precision at *M* = 100. Recall is also improved compared to the arbitrary subnetwork for tMIIC, though only to the random baseline at *M* = 100, 000. For LPCMCI and SVAR-FCI, recall remains low across the two smaller sample sizes tested, due to time limitations for these methods with the GPDCtorch kernel. This improvement in performance of the fast subnetwork relative to the arbitrary subnetwork provides further evidence that homogeneous timescales and stronger memory amplitudes improve the detection of hidden nodes.

## Discussion

In this work we presented a benchmark for causal discovery for the problem of learning networks for biochemical kinetic models from time series data, focussing on reaction kinetics of mass action and Michaelis-Menten type. With knowledge of the ground-truth networks derived from the equations describing the system, we were able to systematically assess the performance of three state-of-the-art temporal causal discovery algorithms: tMIIC, PCMCI+ and its latent variable version LPCMCI, and SVAR-FCI.

To obtain a rigorous random baseline against which to assess performance, we extended the framework for simulation-based negative controls of [42] to this setting of time series causal networks. We compared the inferred networks to randomly generated networks that were consistent (both in terms of sparsity and orientation rules) with the predicted networks they acted as a comparison for. Existing benchmarks for causal discovery typically omit this more rigorous control, and it has been shown that especially for dense networks, it is not difficult for skeleton metrics of precision and recall to come very close or even exceed algorithmic performance [42]. Despite this, we have shown that temporal causal discovery on biochemical kinetic models does outperform these strict random baselines, with tMIIC and PCMCI+ reaching overall higher prediction quality than SVAR-FCI. Furthermore, tMIIC could handle a more substantial amount of samples within pre-set runtime limits and its information-based strategy for conditional independence testing applies seamlessly to linear dynamics as well as nonlinear ones with different functional forms. The advantage of this agnosticism may become particularly evident in real-world scenarios where an a priori guess on the underlying dynamics can prove difficult, or in the presence of highly nonlinear effects – additional benchmarks in this direction would be useful to carry out.

We found that causal discovery can reconstruct the networks under consideration, as long as data sampling regimes (in terms of sampling rates and trajectory length) are consistent with the typical timescales of the variables. We showed on the toy models that, globally, network reconstruction performance improves when sampling rates are chosen with reference average timescales of the system and that node specific performance is optimal when sampling matches the Nyquist rate. Oversampling (sampling more frequently than the timescale) was related to an increase in the prediction of false positive edges, while undersampling (sampling less frequently than the timescale) meant that true edges were not predicted. These results were corroborated by tests on the larger EGFR network, a model with experimentally calibrated parameters. Issues of undersampling can be simply explained as information being lost when sampling occurs more infrequently than the causal interactions, as we could clearly pinpoint by introducing a notion of effective trajectory length (see Eq. 6) and stratifying performance as a function of it. This issue is common when working with real data and several approaches to overcome undersampling have been proposed [45, 46].

We showed that tMIIC and LPCMCI were effective at detecting latent confounders in the toy models through the prediction of bidirected edges. However, truly time-uncorrelated confounders are rare in observational data as almost all physical measurements are influenced by their own past. Detecting hidden intermediate effects through delayed edges was more challenging and we observed that, again, a choice of sampling regimes tailored to the system’s timescales is important. In particular, a priori knowledge of the typical dynamical timescale of the unobserved node, which in practice could be rooted in domain expertise or available characterizations of the biochemical network, proved to be crucial for the correct estimation of the expected delayed effects on the observed subnetwork. These results are consistent with recent work on detecting subnetwork boundary species through temporal inference [14], where trajectory length and sampling intervals tuned to capture all relaxation timescales in the system mattered for accurate performance. When data was oversampled, we detected many false positive delayed edges in the toy models, even without any hidden nodes. However, when sampling rates were optimized for the timescale of the bulk, we were able to detect delayed edges that indicated the presence and timescale of the hidden node that were not predicted when there were no bulk influences on the subnetwork. On the EGFR network, detecting hidden effects through lagged edges was more challenging, though we showed that performance was slightly improved for a subnetwork choice with homogeneous timescales, and stronger memory function amplitudes. Through a comparison to subnetworks dynamically decoupled from their bulk, we found that disentangling the lagged edges present as a result of heterogeneous timescales from those that are due to the presence of hidden nodes is a major difficulty. There is however margin for improvement in terms of identification and characterization of hidden nodes, possibly via the combination of causal discovery to complementary approaches. The less stringent task of detecting boundary nodes (i.e., subject to some bulk influence) can be carried out with high accuracy by targeting the variance of the bulk-induced effective noise through symbolic regression [14], an approach which requires a guess of the basis functions well approximating the subnetwork dynamics and multiple trajectory realizations to estimate the variance. Overall, our findings set quantitative guidelines to help calibrate algorithms’ hyperparameters and design sampling strategies towards an optimal causal reconstruction performance.

One limitation of the causal discovery methods we studied here is their requirement for stationary data (i.e., time series with no trend or seasonal component). Much of the observational data in molecular and cell biology, such as gene regulatory networks through development and cell cycle phases or under changing environmental conditions, are not stationary. However, the core assumption of these methods is of causal stationarity (that causal effects do not change over time) which makes the dynamics amenable to a representation as a time-unfolded network. For this reason, extending temporal causal discovery to a non-stationary setting is challenging. Non-stationary dynamic Bayesian networks (nsDBNs) were introduced by [47]. They relaxed the stationarity assumption by allowing the structure of the network to change at transition times (‘epochs’), learning a specific network structure for each epoch using a score-based approach. In [48], the constraint-based methodology is extended to the non-stationary setting with a method called

CD-NOD (Constraint-based causal Discovery from heterogeneous/NOnstationary Data). This method allows causal relations to change over time by introducing a surrogate variable *C* which is treated as a cause of any variable whose mechanism changes over time. If in the skeleton phase a variable is dependent on *C* given its parents, it is modelled as non-stationary. nsDBNs do not have a mechanism for detecting hidden nodes, while CD-NOD can detect specific types of confounder through bidirected edges. It would be interesting, as future direction, to assess the performance of these methods for biochemical network reconstruction in the presence of hidden confounders and in non-stationary regimes, leveraging the insights we have established here on the interplay between sampling choices and typical evolution timescales.

Other methods have been applied to this problem of learning networks from data. Some such methods aim to directly learn the governing equations of a given system. Sparse Identification of Nonlinear Dynamics (SINDy) [49] learns dynamical systems, similar to those studied here, of the form 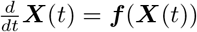 where the function ***f*** is determined from a library of potential functions using sparse regression. There are many extensions to SINDy for equation discovery (e.g. [50–52]) but these are mostly applicable to deterministic setting. Alternative approaches like Approximate Bayesian Computation (ABC) [53, 54] for parameter inference are simulation-based not relying on explicit likelihoods and can be applied to stochastic models easily. A recent approach combines ABC with sparsity inducing priors and Gibbs sampling to develop a general equation discovery approach applicable to stochastic biochemical networks [55]. Other ODE and simulation-based approaches such as hybrid neural models and neural parameter calibration [56–59] iteratively fit parameter values to data using neural networks, aiming to minimize the loss between true and predicted data. These parameter inference and equation discovery methods could be augmented with the causal discovery methods investigated here to constrain the space of possible networks ahead of the parameter learning step.

## Materials and methods

### Numerical solution of SDEs

The equations defining the biochemical systems were numerically solved with the DifferentialEquations.jl package in Julia [60] using an Euler-Maruyama scheme. The SDEs were solved using a time step of 10^−5^ to ensure high numerical accuracy and the trajectories sampled at various values of Δ *τ* : 10^−3^, 10^−2^, 10^−1^. For the more in-depth investigation of timescales and sampling of Fig 5, trajectories were also sampled at Δ *τ* = 10^0^. The larger values of Δ *τ* allow causal discovery algorithms to observe slower dynamics, while the smaller values provide a finer resolution for any fast reactions.

The amplitude of the noise is set by the inverse of the reaction volume *ε* = 1*/V* . We assume the diameter of a mammalian cell is ∼ 20*µm* [61], hence *V* is on the order of (20*µm*)^3^. In the toy models, the mean smallest steady state *y*_*i*_ across the parameter sets is ∼ 0.07, giving an average minimum absolute steady state number of molecules of *V y*_*i*_ ∼ 560. To reproduce this number, we must choose *ε* = 0.0001. For the EGFR model, the smallest steady *y*_*i*_ *∼* 0.05, making the smallest copy number *V y*_*i*_ *∼* 400 and so also requiring the choice of *ε* = 0.0001.

Code is available at https://github.com/holly-chambers/biochemical-kinetic-causal-discovery.

### Coarse-grained SDEs with hidden nodes

In the case of partial observation of the biochemical system, we treat the ‘observed’ species (i.e., whose measurements are available) as forming a subnetwork embedded within a larger, unobserved network called the bulk, according to the framework of [10–12].

### Subnetwork-reduced linearized dynamics

In the case of the linearized dynamics, the one we studied here in connection to hidden nodes, it is straightforward to derive the subnetwork-reduced dynamics accounting for the dynamical effect of the hidden nodes (i.e., the bulk). Following [11], the *N* -dimensional vector of concentration deviations *δ****X***(*t*) is split into *N* ^s^ subnetwork species stored in the vector *δ****X***^s^, and the *N* ^b^ bulk species in the vector *δ****X***^b^.

Correspondingly, the linearized rate matrix ***K*** of Eqs. 3 and 4 is decomposed into a block form as shown in Eq 7, where ***K***^ss^ and ***K***^bb^ represent reactions between purely subnetwork or bulk species respectively, and ***K***^bs^ and ***K***^sb^ reactions of involving the subnetwork’s boundary species (i.e., reactions between subnetwork and bulk species). Eq 3 may then be decomposed into equations for the subnetwork and bulk species as

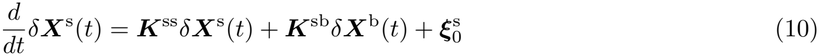

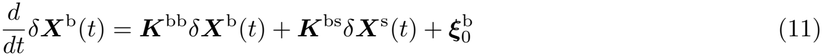

with the covariance matrix of Eq 5 decomposed similarly as

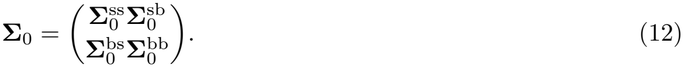

To obtain the dynamics in terms of only the subnetwork, we first solve for *δ****X***^b^(*t*) with initial condition δX^b^(0)

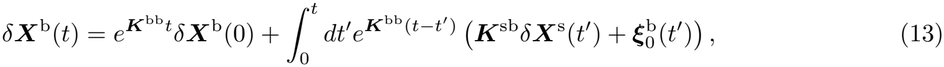

and then substitute into Eq 10. One obtains

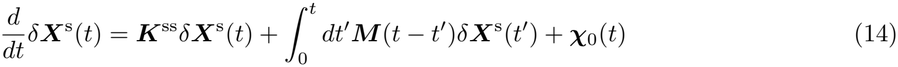

where the memory function ***M*** (*t− t*^*’*^), accounting for the influence of the subnetwork on the bulk fed back into the subnetwork at a later time, is given by Eq 8 and

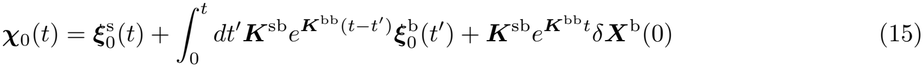

is an effective coloured noise incorporating intrinsic noise from the subnetwork 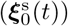, the effect of past intrinsic noise from the bulk 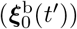 at time *t*^*’*^ on the subnetwork at time *t*, and the initial condition of the bulk deviations (*δ****X***^b^(0)). The time correlation of the effective coloured noise ***x***_0_(*t*) is given by

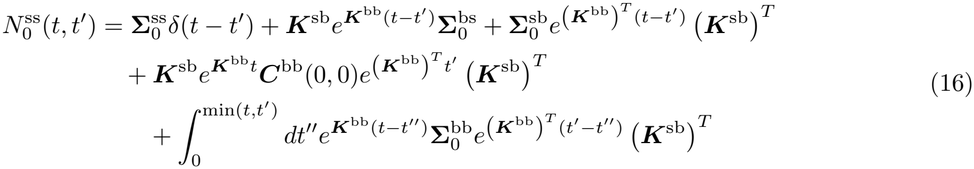

where ***C***(*t, t*) is the equal time correlation matrix which obeys

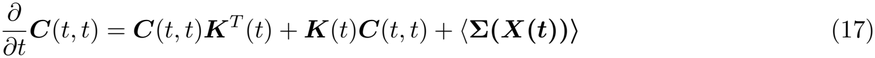

then to find ***C***(0, 0), we set 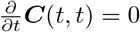 and solve the Lyapunov equation

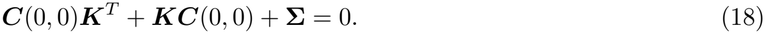

In the case of a single-species bulk with steady-state initial conditions, such as the one studied on the toy model, 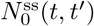 can be simplified as

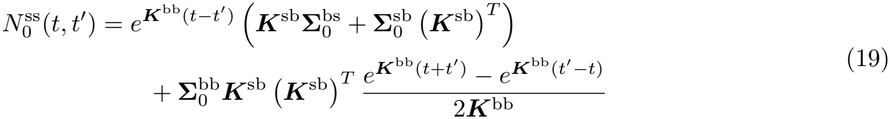

Overall, from Eq 14, one can see that the pure subnetwork reactions appear in their original form through the matrix ***K***^ss^, while the effect of the bulk induces non-Markovian effects through the memory function and time-correlated (coloured) noise.

#### Timescales of non-Markovian terms

The amplitude of the memory function is given by ***M*** (*t − t*^*’*^) from Eq 8 evaluated at zero time difference, *t −t*^*’*^ = 0 and is simply ***M*** (0) = ***K***^sb^***K***^bs^, as at *t t*^*’*^ = 0 the exponential reduces to the identity matrix. The timescale of the decay of a generic memory function ***M*** (*t*) can be derived, following [10], as its time integral divided by its amplitude

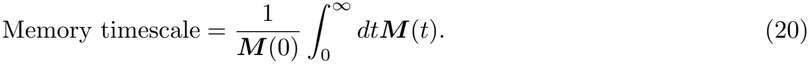

In this way, if ***M*** (*t*) decays as an exponential, 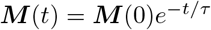, then Eq 20 gives the actual decay time *τ*. Applying this to the memory function of Eq 8 and using ***M*** (0) = ***K***^sb^***K***^bs^ as derived above gives the self-memory timescale of Eq 9.

For the Michaelis-Menten toy model with the *X*_3_ node hidden in the bulk and the remaining nodes in the subnetwork, ***M*** (0) is given by

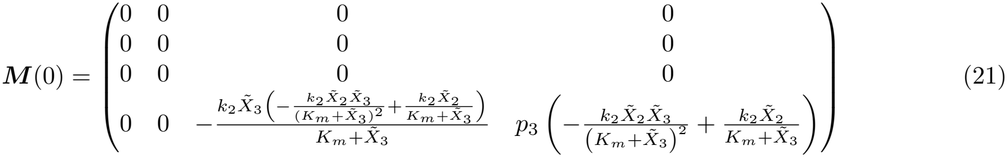

where 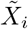 is the steady state of species *i*. Here, the variables are ordered as [*X*_1_, *X*_1*p*_, *X*_2_, *X*_3*p*_] and the sparse structure of this matrix indicates that there are memory effects acting only on *X*_3*p*_ (the final row of the matrix) and these come from *X*_2_ and *X*_3*p*_ itself. We can compute the timescale of the self-memory of *X*_3*p*_ on itself according to Eq 20 as

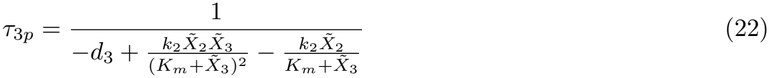

which is equivalent to 1*/K*^bb^ in this single-species bulk case.

### Computing timescales of biochemical systems

We calculated timescales of variables in the biochemical reaction systems by assuming that the autocorrelation function (ACF) of the time series decays exponentially. We computed these autocorrelations for each variable from their trajectories, and fit an exponential decay model with rate of 1*/t*^*\**^ (where *t*^*\**^ is the timescale for the variable) using curve fit from scipy.optimize [62]. The characteristic timescale 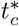 was set to the slowest timescale across the species in the system.

We computed the coefficient of variation (CV) of the timescales to measure the variability between them. This is the ratio of the standard deviation to the mean of the timescales.

For the analysis of Fig 5, we chose parameter sets representing fast/slow and homogeneous/heterogeneous timescales according to the two measures described above. Note that the slow homogeneous parameter set was chosen in such a way as to ensure that the timescale of *X*_2_ was sufficiently slow (of order 1) for the comparison, as it is connected to all other variables and strongly influences reconstruction performance, hence the choice to not select the point in the very bottom right corner of Fig 5B.

### Deriving causal networks from SDEs

We considered our system of equations (1) as structural causal kinetic models [63], i.e., the time evolution of each variable is a function of its causal parents

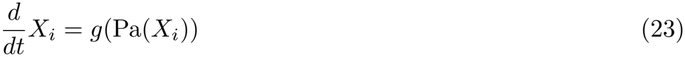

In this way, we constructed the ground truth causal network by directing an edge from 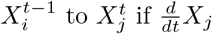 depends on *X*_*i*_ (i.e., if *X*_*i*_ appears in the dynamic equation for *X*_*j*_). This is equivalent to constructing the adjacency matrix of the summary causal network by setting any non-zero elements of the Jacobian matrix (equivalent to the matrix ***K*** in this work) to 1. Due to the Markovian nature of the equations, we consider each of these effects to occur with a Δ*t* = 1.

### Causal networks with hidden nodes

When hidden nodes are present, the corresponding causal subnetwork is represented as a time series Maximal Ancestral Graph (ts-MAG). Only the subnetwork variables that have a hidden node as a direct cause (i.e., boundary nodes) will receive additional edges in the ts-MAG.

The nature of these additional edges depends on the role of the hidden variable. For instance, if the hidden variable is a time-uncorrelated confounder (as in the augmented toy models of Fig 6 and S3 Fig), it induces bidirected contemporaneous edges between the boundary nodes. Conversely, if the hidden variable has autocorrelations, various types of time-delayed and contemporaneous edges may be induced. These additional edges of the ts-MAG can be inferred from the topology of the summary network of the ts-DAG alone (we provide derivations and proofs in S1 Appendix).

When a hidden node causes a boundary node, additional auto-link edges with 1 *<* Δ*t* ≤_max_ are induced. We see these in our toy models and the EGFR network with hidden nodes, any time a boundary node is present. When a hidden node acts as an intermediate between two subnetwork nodes, the boundary node will acquire delayed cross-link edges for all 1 *<* Δ*t* ≤ _max_. In the toy model of Fig 6, *X*_3_ is an intermediate *X*_2_ → *X*_3_ → *X*_3*p*_, and in the ts-MAG, delayed edges between *X*_2_ → *X*_3*p*_ are induced. For the case where the hidden node is an autocorrelated confounder, each of the boundary nodes receive cross-link edges from each of the other boundary nodes, for all Δ*t ≤*_max_. We show examples of autocorrelated confounders in the EGFR network, and in the supplementary example of the mass action network with hidden nodes S3 Fig. The final type of edge induced in a ts-MAG is when a boundary node has other causes, aside from the hidden node. In this case, delayed edges are present from these other causes into the boundary node for all 1 *<* Δ*t ≤*_max_. This type is exemplified by the delayed cross-link edge from *X*_1_ to *X*_2_ in S3 Fig.

Putting these results together, one can derive the block adjacency matrix ***Ã*** of the time-unfolded subnetwork-reduced ts-MAG from the structure of the reaction rate matrix, Eq 7 (for a derivation starting from the adjacency matrix of the time-unfolded full-system causal graph see S1 Appendix). Each block is of size *N* ^s^ × *N* ^s^ and a single block ***Ã***_*t,t−*Δ*t*_ represents links from time *t* − Δ*t* to *t*. The blocks Δ*t* = 0, Δ*t* = 1, and 1 *<* Δ*t* ≤ _max_ are each defined in a distinct way and are given by

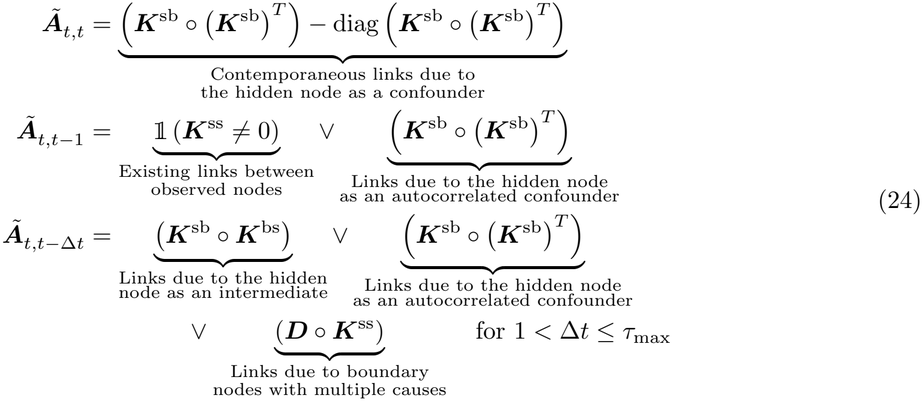

where 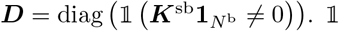. is an indicator function, 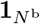 is the vector of *N* ^b^ ones so that ***D*** is an *N* ^s^ × *N* ^s^ diagonal matrix with ***D***^*ii*^ = 1 if and only if *X*^*i*^ is a boundary node. ∨ is the logical ‘or’ operation and the boolean matrix operation denoted by is defined as (***U*** ○ ***V***)_*ij*_ = ∨ _*k*_ ((*U*_*ik*_ ≠ 0) ∧ (*V*_*kj*_ ≠ 0)) with ∧ being the logical ‘and’.

### Network inference with causal discovery

The trajectories generated via numerical solution of the SDE system of interest were passed to the causal discovery algorithms described below. For experiments with hidden nodes, the trajectories for the variables to be hidden were omitted from the data inputted to the algorithms.

The three algorithms benchmarked here are constraint-based causal discovery methods, built upon methods for learning static (non-temporal) causal graphs. The PC (Peter and Clark) [18] algorithm is the foundational constraint-based algorithm for static data with no hidden nodes. Starting from a complete graph, it iteratively removes edges between variables that are conditionally independent given some subset of the remaining variables, before orienting edges by identifying v-structures and applying acyclicity constraints. The Fast Causal Inference (FCI) algorithm [19] extends PC to allow for latent confounders, indicating them with bidirected edges (*X Y*) between the variables on which they act. The three algorithms considered here extend this to time series data each accounting for autocorrelation, contemporaneous effects, and latent variables as described in the following subsections.

### tMIIC

MIIC (Multivariate Information-based Inductive Causation) [64] and its temporal extension tMIIC [23] use information theory to construct causal networks. The algorithm begins with a fully connected graph of all variables and iteratively removes dispensable edges when independence is found using conditional multivariate information *I* (e.g., an edge between *X* and *Y* is removed when *I*(*X*; *Y* | {*A*_*i*_}) = 0, where {*A*_*i*_} is the separating set). The undirected skeleton graph is then oriented, first using that negative 3-point information implies the existence of a v-structure [65]. These v-structure orientations may then be propagated downstream following the rules of ancestral graphs [66].

tMIIC extends the static version of MIIC to construct time-unfolded networks (as in Box 1), using the same procedure for the skeleton discovery, but for orientation first directing any delayed edges forward in time before applying v-structure rules on contemporaneous edges.

The trajectories were passed to tMIIC via the MIIC package in R [23, 64]. We used the time series mode of MIIC (tMIIC, v2.0.3), specifying that there are two layers (*τ*_max_ = 1) and setting the hyperparameter Δ*t* = 1 so that we have only one time step between each layer, consistent with the Markovian nature of the dynamics. All other hyperparameters for the algorithm use the default settings: mode = ‘TS’ (time series mode of MIIC); latent = no; cplx = nml (due to the finite input size, corrections to the information measures are made by shifting by normalized maximum likelihood); orientation = TRUE (edges are assigned directions); ori proba ratio = 1 (all orientations are accepted no matter probability of orientation); propagation = FALSE (contemporaneous edges are only oriented with v-structure orientations); conf threshold = 0 (no threshold to filter less probable edges following the skeleton step). We chose not to filter edges at this step, as the method typically underestimated the sparsity of the networks.

For experiments with hidden nodes, the trajectories for the hidden nodes are omitted from the input data and the tMIIC input argument indicating that latent variables should be accounted for was set to ‘true’. In this setting, we also chose the number of layers adaptively, according to 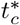 so that delayed edges indicating the presence of hidden nodes could be detected. For the analysis of hidden nodes on the toy models of Fig 6 and S3 Fig we used 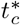 of the subnetwork, while for the EGFR model (Fig 7), we compared the effect of choosing 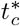 of the subnetwork to 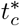 of the bulk.

### PCMCI+ and LPCMCI

PCMCI (Peter and Clark Momentary Conditional Independence) [67] extends the PC algorithm to time series data following the approach explained in Box 1. The edge removal by conditional independence tests step is split into two parts: a selection phase (PC_1_) and a testing phase (MCI). In PC_1_, irrelevant variables are iteratively removed by testing for conditional independence with increasingly complex conditioning sets building up to test only the strongest associations. Having reduced the conditioning set to a much lower dimensionality, the momentary conditional independence (MCI) test assesses if a link exists by testing the conditional independence between a lagged source variable and its contemporaneous target given additionally the lagged parents. This accounts for inherent autocorrelations in time series data, controlling for false positives.

PCMCI+ [20] extends this method to also learn contemporaneous edges. In this approach, lagged and contemporaneous edges are removed in separate phases to reduce the number of conditional tests to be performed.

LPCMCI [21] is a further extension of the above two methods, to handle latent confounders. This method combines the edge removal and orientation phases to learn ancestral relations early on. This iterative process discards conditioning sets that contain known non-ancestors of the variables being tested, and uses known causal parents (identified in the iterative learning process) as default conditions for subsequent conditional independence tests. This approach is designed to increase the effect size of conditional independence tests leading to a greater recall of true links.

We used PCMCI+ to learn networks for fully observed data, and LPCMCI to learn networks with hidden nodes from partial observations through the Tigramite Python package [22]. For nonlinear data, we used the GPDCtorch (Gaussian process regression and distance correlation) kernel to test for conditional independence, and for data from the linearized models the ParCorr (partial correlation) kernel. We set *τ*_max_ = 1 with fully observed data, and chose *τ*_max_ adaptively for experiments with hidden nodes. The remaining hyperparameters were left at default settings: link assumptions = None (no assumptions made about which variables do/do not have an edge); pc_alpha = 0.01 (significance level of conditional independence test); auto_first = True (autodependency links are tested first). Additionally for LPCMCI n_preliminary iterations = 4 (using the maximum number of iterations in the preliminary phase of the algorithm).

### SVAR-FCI

SVAR-FCI (Structural Vector Autoregression Fast Causal Inference) [24] is the temporal extension of the FCI algorithm described above. Time ordering is respected by restricting conditioning sets between variables to only the past or present, and enforcing orientations forwards in time. Like the other constraint-based methods described, causal stationarity is assumed and the learnt network has a repeating structure within a window of size *τ*_max_. Consistent with tMIIC and LPCMCI, SVAR-FCI also allows for both contemporaneous causes and latent confounders acting on observed variables.

The SVAR-FCI algorithm was also run through the Tigramite package in Python with *τ*_max_ = 1 for predictions on the full network, and chose *τ*_max_ adaptively when considering hidden nodes. For consistency with the PCMCI-like methods, pc alpha was again set to 0.01.

### Analysis of performance

The networks inferred by the causal discovery methods were compared to the ground-truth networks. The numbers of true positives (TP), true negative (TN), false positives (FP), false negatives (FN), and misoriented edges (MO) were counted to build the contingency matrix as shown in Fig 2. We classify an edge as MO when it is correctly identified to be present, but has been assigned an incorrect (or no) direction.

Only when the edge is both identified and assigned with the correct direction it is counted as TP. From these, precision, recall and F-score can be calculated for the skeleton (skel.), and fully oriented network (ori.). These metrics are defined in the following way:

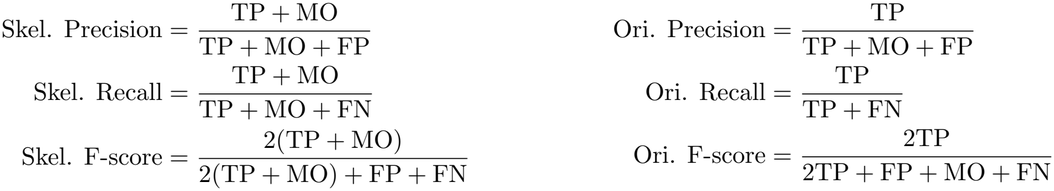

We refer to the stricter, orientation-aware metrics simply as precision, recall and F-score in the text. These metrics may be considered in a time-unfolded or time-collapsed sense. Time-unfolded metrics require the lag of the predicted edge to match the ground-truth lag to be classified as a TP (otherwise the edge is an FP). For time-collapsed metrics, the lag is ignored and a TP is counted as any edge between two nodes that exists in the ground truth.

We also computed performance metrics at the node level, defining node-specific precision, recall and F-scores by counting the numbers of TP, TN, FP, FN and MO for edges to a specific node and calculating precision, recall and F-score as above.

When considering performance accounting for hidden nodes, we focussed on the identification of edges that are typically indicative of the presence of hidden nodes: instantaneous and delayed edges. As such, precision and recall are computed over this set of edges, and TPs are the additional edges induced by the hidden nodes as discussed previously. Delayed edges found in the ground truth may have any lag ≥2 to be considered as a TP.

### Comparison to a random baseline

We computed random baselines to act as a negative control according to the methodology of simulation-based negative controls from [42], extended to a time-unfolded network setting.

For each network predicted by a causal discovery algorithm, we generated 100 random networks, matching the sparsity of the given prediction and redistributing the edges uniformly at random amongst the nodes. These random networks were generated in such a way as to be consistent with the properties of time-unfolded networks, as explained in what follows. Sparsity over the instantaneous, Markovian and delayed edges (i.e., lag=0, lag=1, and lag≥1, respectively) was computed separately in such a way as to force the random networks to have a sparsity level consistent with the prediction for each edge type. For the instantaneous layer, edges were first oriented as directed and then, following the convention of [42], this part of the network converted to a CPDAG using the pcalg package in R [68] for consistency with the causal discovery methods we made comparisons against. For the Markovian and delayed regions, after computing the sparsity of the prediction, edges were randomly redistributed over the same set of layers. Edges with lag ≥1 must always be directed into the future (→), hence all delayed edges were automatically oriented as such.

Performance metrics were computed for these random networks in the same way as if the network had been predicted by causal discovery, that is, against the same ground truth.

## Supporting information

**S1 Table Average F-score increase over random baseline**.

**S1 Fig Overview of the performance of three causal discovery methods for biochemical network reconstruction – linearized dynamics**.

**S2 Fig Performance of network reconstruction by tMIIC with changes to sampling interval of the data – Michaelis-Menten model**.

**S3 Fig Summary of additional edges predicted with hidden nodes – linearized mass action model**.

**S4 Fig Choosing the optimal sampling for the EGFR network**.

**S5 Fig Results of causal discovery on the EGFR network – linearized dynamics**.

**S6 Fig Memory amplitudes and timescale changes in the EGFR networks with hidden nodes**.

**S1 Appendix. Derivations and proofs for identifying additional edges induced by hidden nodes in temporal causal networks**.

## Acknowledgments

The authors acknowledge useful discussions with Nikita Lagrange, Franck Simon, Tiziana Tocci, Louise Dupuis, Orianne Debeaupuis, Mauricio Barahona and Omer Karin.

## Supplementary figures

**S1 Table.** Average F-score increase over the random baseline. Sample sizes (*M*) are 1000 for nonlinear and 100,000 for linearized data unless otherwise specified.

| Data Type | Method | Mass Action | Michaelis-Menten | EGFR |
|---|---|---|---|---|
| **Nonlinear** | tMIIC (*M* = 1000) | +0.2313 | +0.2445 | +0.2038 |
|  | tMIIC (*M* = 100,000) | +0.3326 | +0.3462 | +0.2296 |
|  | PCMCI+ (GPDCtorch) | +0.2990 | +0.2658 | +0.1860 |
|  | SVAR-FCI (GPDCtorch) | +0.2338 | +0.1919 | +0.1189<sup>†</sup> |
| **Linear** | tMIIC | +0.3115 | +0.3440 | +0.2387 |
|  | PCMCI+ (ParCorr) | +0.3421 | +0.3637 | +0.0738 |
|  | SVAR-FCI (ParCorr) | +0.0804 | +0.1610 | +0.1236<sup>†</sup> |

<sup>†</sup> SVAR-FCI results for EGFR use *M* = 100 due to computational limitations.

**S1 Fig.**
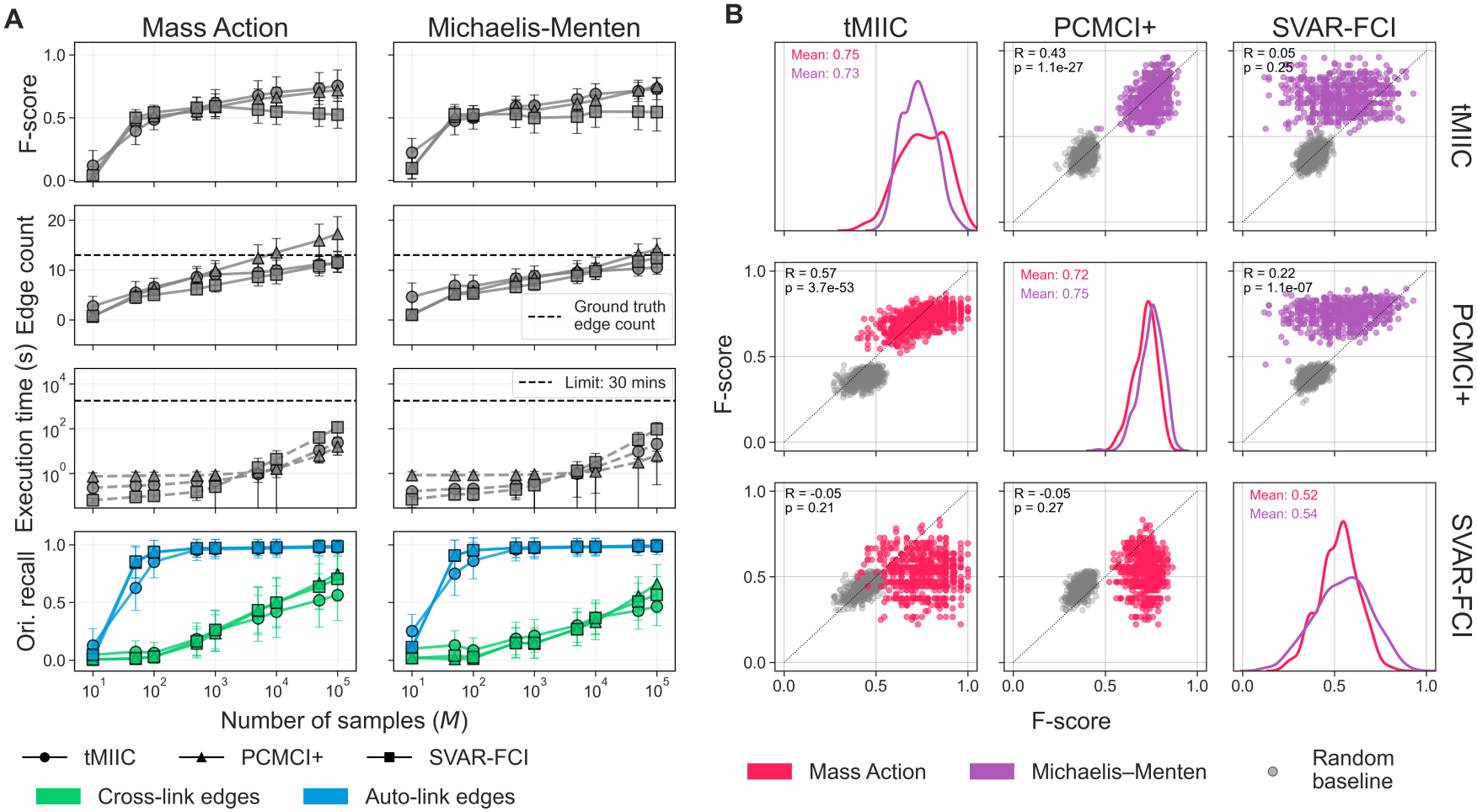
Performance of three causal discovery methods for biochemical network reconstruction for the five-species toy models – linearized dynamics. Linearized mass action and Michaelis-Menten models, Δτ = 0.1. PCMCI+ and SVAR-FCI methods use the ParCorr kernel. A) Mean F-score, edge count, execution time and orientation recall against number of samples *M*, across networks inferred from 600 parameter sets. B) Pairplot of F-scores for the three methods (*M* = 100, 000) across 600 parameter sets and both models. Diagonals show F-score density per method and model, off-diagonals show pairwise scatter with Pearson’s *R* and corresponding p-value.

**S2 Fig.**
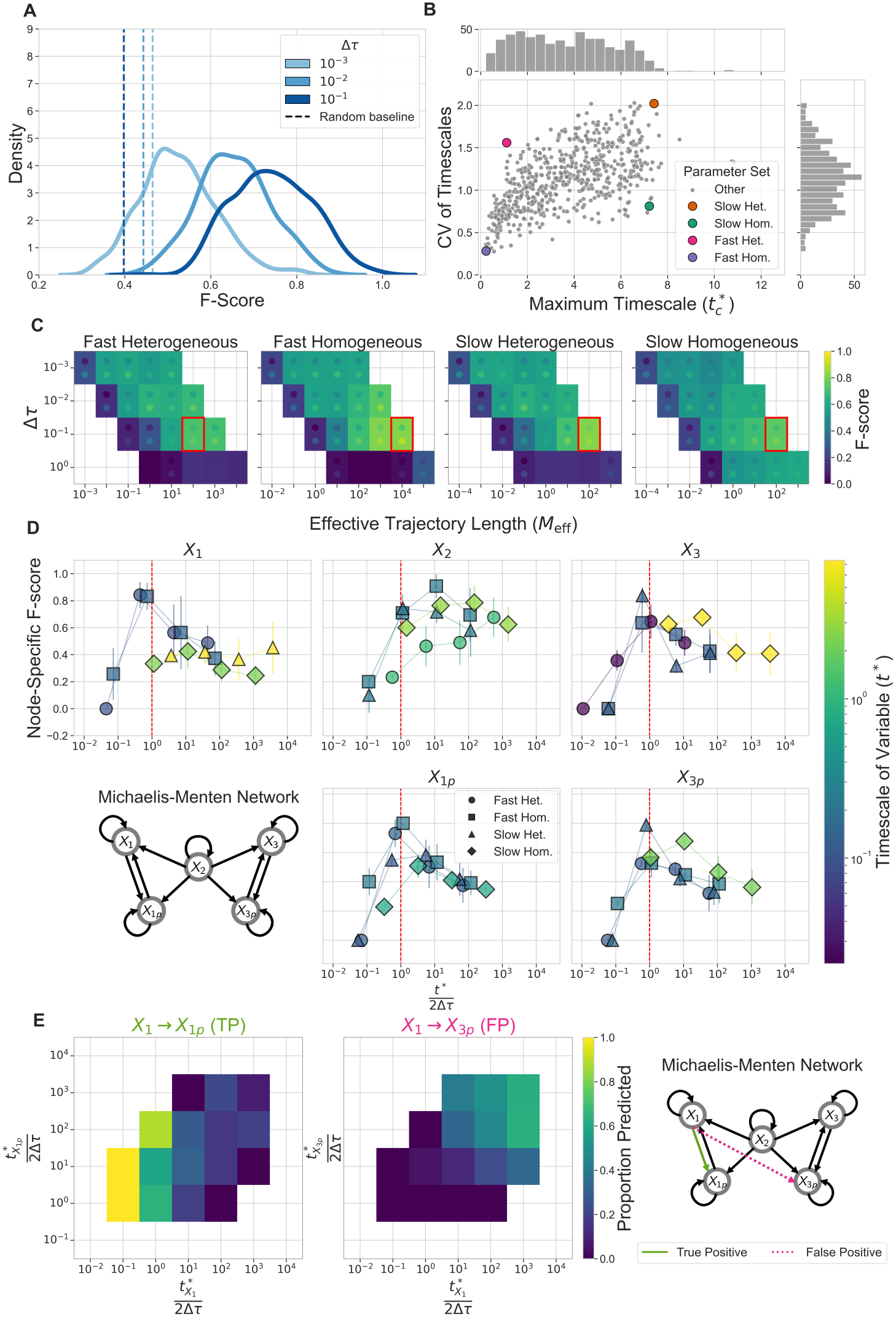
Performance of network reconstruction by tMIIC with changes to sampling interval of the data – Michaelis-Menten model. Nonlinear Michaelis-Menten models, *M* = 100, 000. A) F-score density across 600 parameter sets at three values of Δτ. B) Characteristic timescale (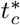, maximum relaxation time across the five timescales) against timescale heterogeneity (coefficient of variation, CV) for each parameter set, with four selected parameter regimes (fast/slow, homogeneous/heterogeneous) highlighted. C) Mean F-score heatmaps for the four selected regimes, varying *M*_eff_ and Δτ, averaged over 10 realizations. Dots mark ±1 SD. D) Node-specific F-scores against *t*^*\**^*/*2Δτ for each variable. Red dashed line shows sampling interval equal to Nyquist rate (*t*^*\**^*/*2Δτ = 1). E) Proportion of predicted edges as a function of *t*^*\**^*/*2Δτ for both variables in a representative true positive edge (left), and representative false positive edge (right), across all 600 parameter sets and three values Δτ = 10^−3^, 10^−2^, 10^−1^ (1800 networks total). Bins with *<* 10 observations omitted (1 excluded for the true positive edge, 1 for the false positive edge).

**S3 Fig.**
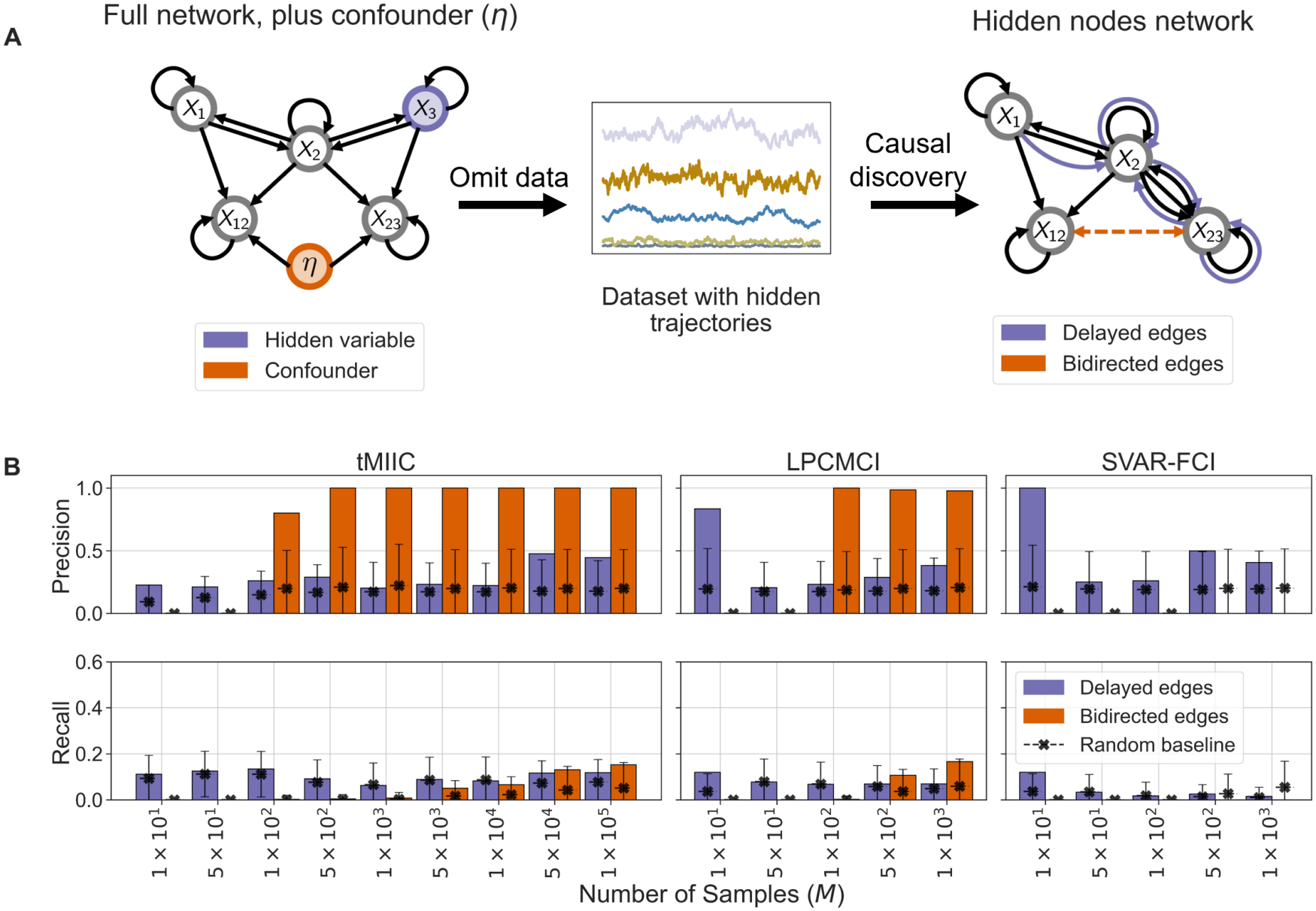
Summary of additional edges predicted with hidden nodes – mass action model. A) Full network with uncorrelated confounder *η* (orange), and hidden variable *X*_3_ (purple). Hidden trajectories are withheld from causal discovery (centre), to obtain the reduced causal network (right). The hidden node induces delayed edges (purple) and bidirected edges (orange) are induced by the confounder. B) Precision and recall of the delayed and bidirected edges indicating hidden nodes across sample sizes *M*, for tMIIC, LPCMCI and SVAR-FCI (nonlinear data, Δ*τ* = 0.1, *τ*_max_ set according to the slowest timescale in the subnetwork, GPDCtorch kernel for LPCMCI and SVAR-FCI, maximum *M* limited by 30-minute limit for computation). Black crosses mark random baseline values.

**S4 Fig.**
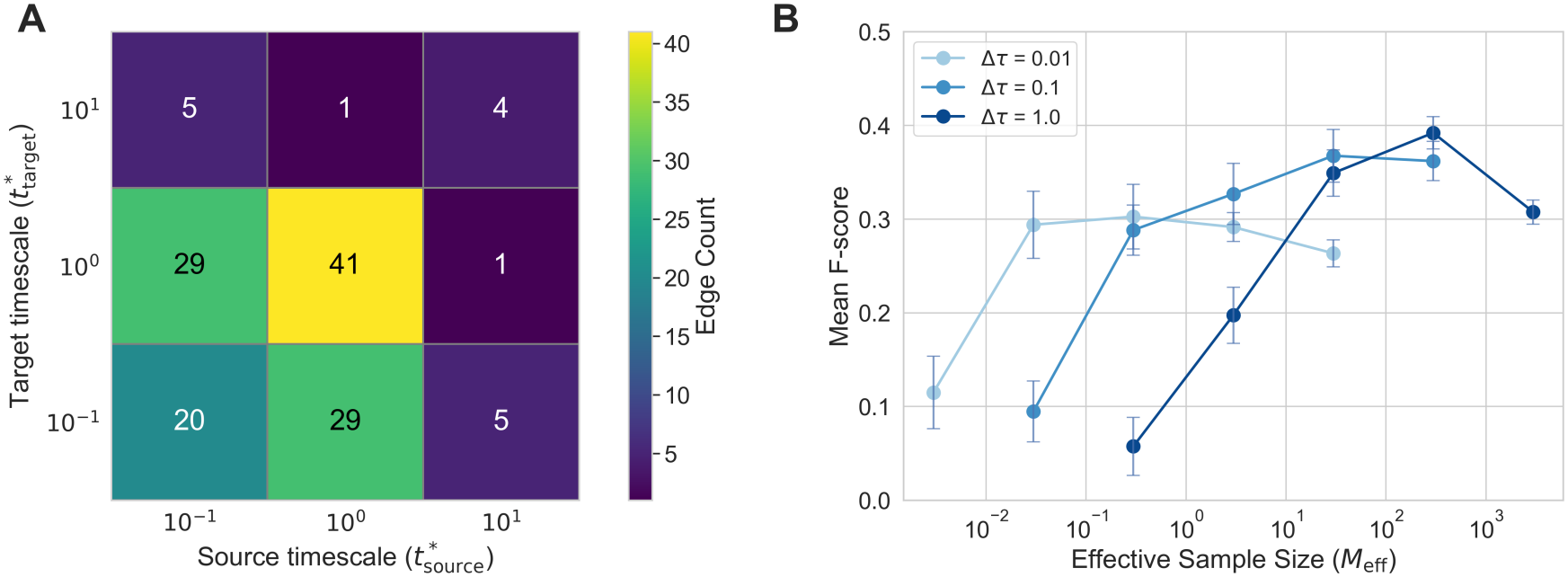
Choosing the optimal sampling for the EGFR network. A) Heatmap summarising the timescales of each pair of source-target variables of the EGFR ground-truth edges. B) Line plot for the mean time-collapsed F-score over 10 seeds of the nonlinear EGFR model as a function of the effective trajectory length *M*_eff_ for three values of Δ *τ* . Error bars show the standard deviation.

**S5 Fig.**
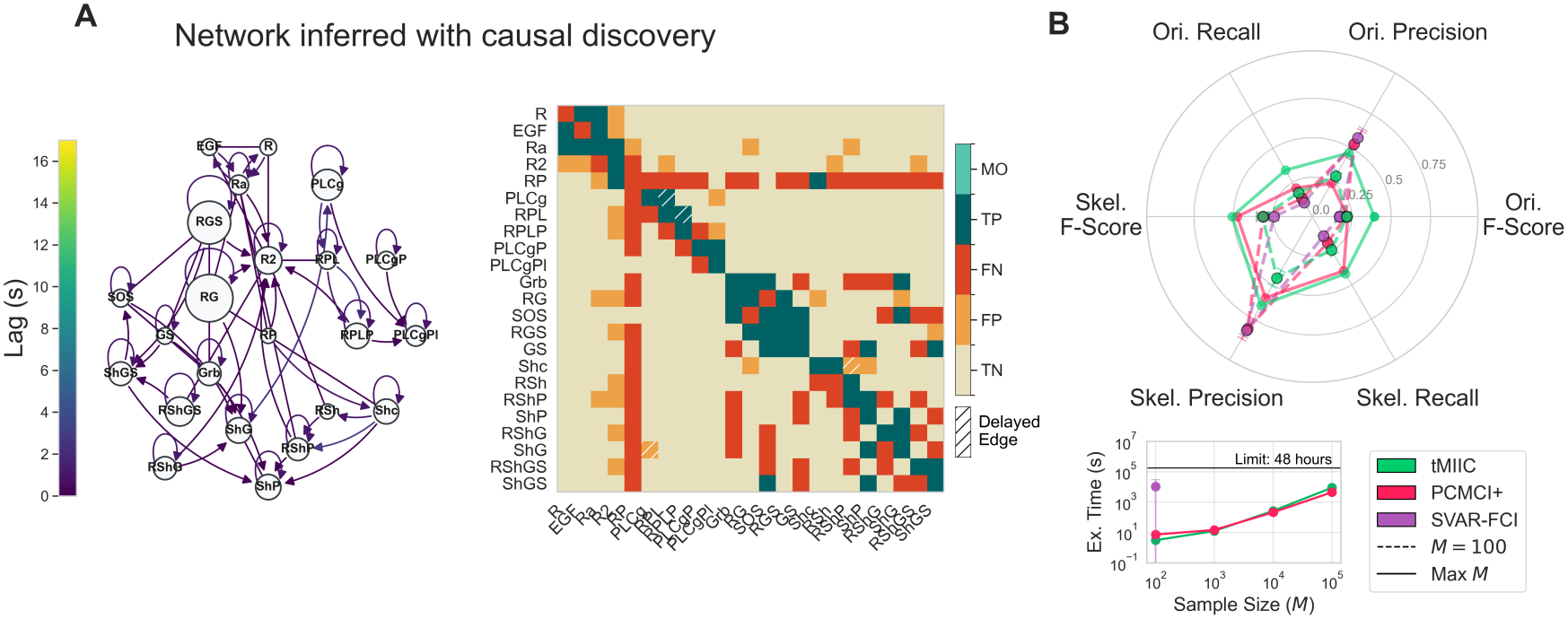
Results of causal discovery on the EGFR network – linearized dynamics. A) Example network and time-collapsed heatmap representation learnt by tMIIC. Linearized model, Δ *τ* = 1, *τ*_max_ = 17, *M* = 100, 000. B) Radar plot comparing the three methods’ performance. Dashed lines show performance metrics at *M* = 100, the maximum common sample size, while solid lines show performance at maximum *M* = − 100, 000 for PCMCI+ and tMIIC. Inset shows execution time against sample size. Linearized data, Δ *τ* = 1, *τ*_max_ = 17, ParCorr kernel for PCMCI+ and SVAR-FCI.

**S6 Fig.**
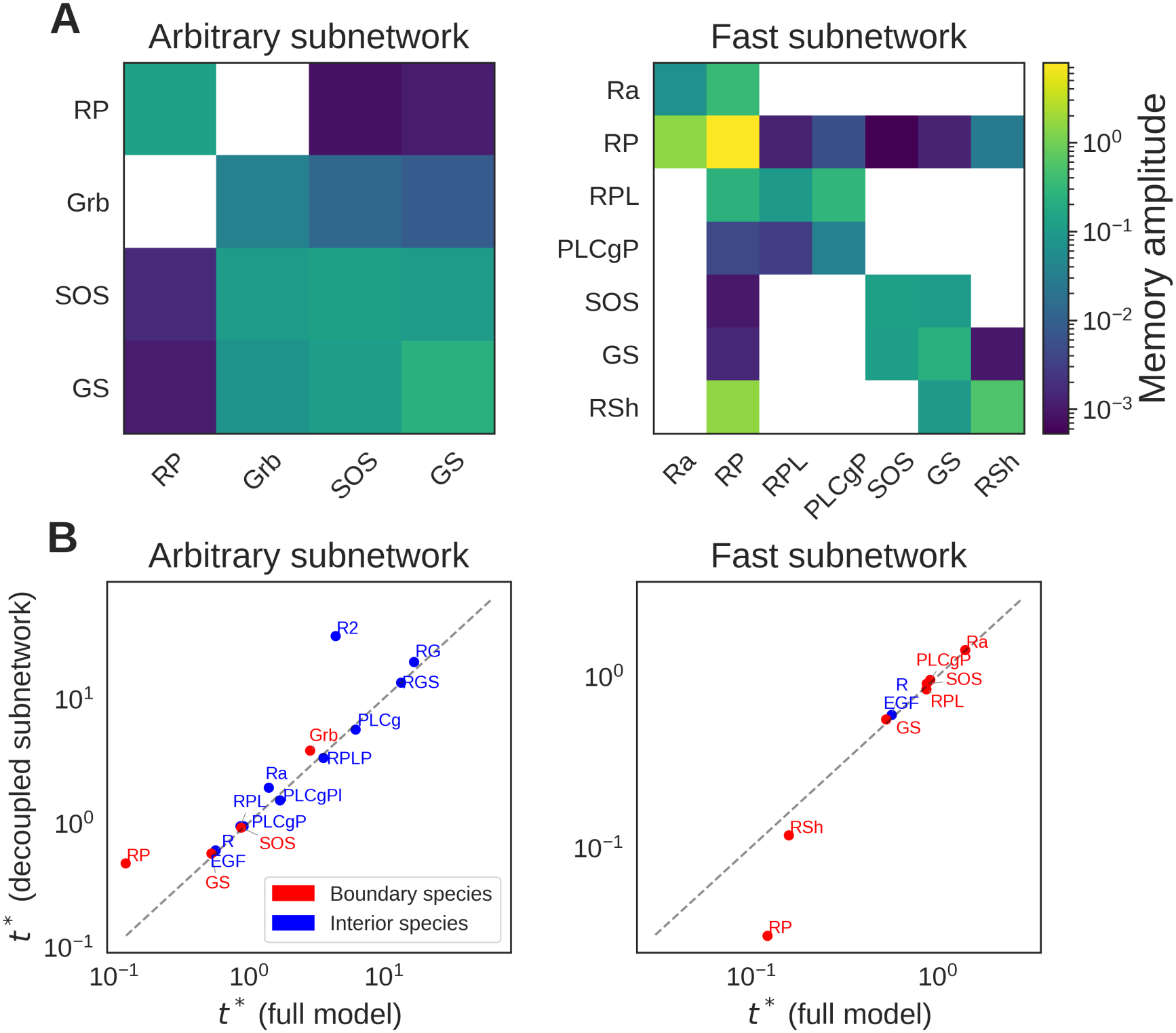
Memory amplitudes and variable timescales in the two subnetworks. A) Heatmaps of memory amplitudes of boundary species in the arbitrary subnetwork (left) and fast subnetwork (right). B) Comparison of timescales in the full and decoupled models in the arbitrary subnetwork (left) and fast subnetwork (right).

## Supplementary text

### 1 Setup

#### Time series Directed Acyclic Graph

We consider a multivariate time series {***X***^*t*^}_*t∈* ℤ_ with *N* variables, generated by a first order vector autoregressive process

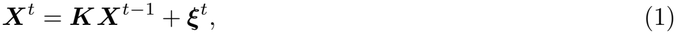

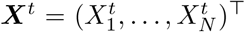 is the vector of variables at time *t*, ***K*** ∈ ℝ ^*N × N*^ is the coefficient matrix, and ***ξ*** is an *N* -dimensional zero-mean Gaussian white noise process.

Eq. 1 has a causal interpretation through its associated time series directed acyclic graph (ts-DAG) such that there exists a directed lag 1 edge 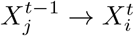 if and only if ***K***_*ij*_ ≠ 0.

Throughout, we make the following assumptions:

##### Assumption 1.

*The process is first-order and time-homogeneous*.

##### Assumption 2.

*There are no contemporaneous directed edges*.

##### Assumption 3.

*All variables have a causal influence on themselves, i*.*e*., ***K***^ii^ ≠ 0 *for all i*.

##### Assumption 4.

*The process is stable (i*.*e*., *all eigenvalues of* ***K*** *are real and strictly negative), so that the solution to Eq. 1 is stationary*.

As explained in Box 1 in the main text, to represent the time-unfolded ts-DAG, we use a block adjacency matrix whose rows correspond to target nodes, and columns to source nodes. We order the nodes, firstly by variable and then by lag (i.e. 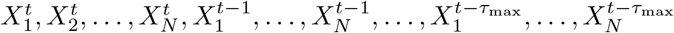) enforcing a block structure in the matrix, with (for example) ***A*** _*t,t*− 1_ denoting links from nodes at *t* − 1 to nodes at *t*.

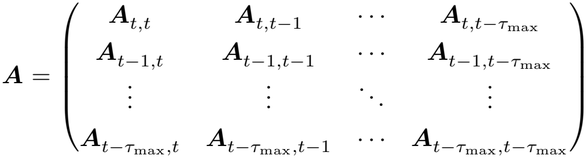

Due to Assumption 1, the process is first-order and so described by the directed links from time *t* − 1 to time *t*. Furthermore, the time-homogeneity corresponds to causal stationarity and so in fact the block ***A***_*t,t−*1_ provides the full description, hence ***A***_*t,t −* Δ*t*_ = **0** for all Δ*t >* 1. Additionally, by Assumption 2, there are no contemporaneous directed links and so ***A***_*t,t*_ = **0**.

The adjacency matrix corresponding to the system of Eq. 1 can be derived from the matrix ***K*** . The top right block, ***A***_*t,t −* 1_ = (***K*** ≠ 0) = 1_***K***_ where 1 is the indicator function transforming all non-zero elements of ***K*** to one. The bottom left block is the transpose of this, 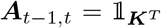. Since we assume that the process is Markovian and first order, we require only these two blocks in the adjacency matrix, i.e.,

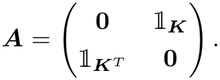

#### Time series Maximal Ancestral Graph

If we assume that *N* ^b^ time series components (the ‘bulk’) are unobserved (or hidden) and the *N* ^s^ = *N* − *N* ^b^ subnetwork components remain observed, then we may partition ***X***^*t*^ into observed (s) and unobserved (b) components as

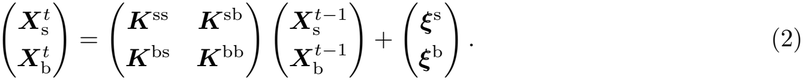

***K*** is also partitioned with ***K***^ss^ and ***K***^bb^ describing links between purely observed and purely unobserved nodes, respectively, while ***K***^sb^ describes the influence of the unobserved processes on the observed and ***K***^bs^ the observed processes on the unobserved. Similarly, each block of the adjacency matrix ***A***_*t−* Δ *t,t−* Δ*t*_ can be partitioned to describe causal links between each combination of observed and unobserved nodes.

The corresponding causal network obtained after marginalising out the hidden variables is the time series maximal ancestral graph (ts-MAG). Here, the focus is on the adjacency structure of the observed time-unfolded ts-MAG. We characterise which observed node pairs become adjacent after integrating out the hidden variables, and at what lags these adjacencies occur. We assume throughout that there are no selection variables acting on the system:

##### Assumption 5.

*There is no selection bias (the set of selection variables is empty)*.

### 2 Identification of hidden node-induced edges

To find the hidden node-induced additional edges in a ts-MAG, one needs to identify the so-called *inducing paths* between the observed variables with respect to the hidden nodes.

#### Definition

(Inducing path [1]). An inducing path is a path between two observed nodes *X*_*i*_, *X*_*j*_ on which every non-endpoint node is either in the set of hidden nodes, or is a collider, and every collider is an ancestor of either *X*_*i*_ or *X*_*j*_.

Additional observed adjacencies appear in the ts-MAG if an inducing path can be found between them in the ts-DAG with respect to the set of hidden nodes [2].

We now present some results for systematically identifying the additional edges in a ts-MAG due to the presence of hidden nodes. We characterise these edges according to the type of hidden node from which they are derived and show that additional edges may arise from three key types of interaction with boundary nodes: hidden nodes as confounders; hidden nodes as intermediates; and boundary nodes with multiple causes. These results are based on the inducing path construction introduced above and are all consequences of the following proposition.

We will call an observed node a *boundary node* if it has at least one hidden parent in the ts-DAG, and index as *a, b, c*. We will call all other observed nodes *interior nodes* and index as *i, j, k*. We will call nodes for variables of unobserved components *hidden nodes* and index as *l, m, n*.

#### Proposition 1.

*Only variables with hidden nodes as a cause in the ts-DAG will receive additional edges in the ts-MAG*.

*Proof of Proposition 1*. Nodes receive additional edges in a ts-MAG if an inducing path can be found between two remaining observed nodes with respect to the set of hidden nodes. By the definition of an inducing path, the non-endpoint nodes on the path may only be in the set of hidden nodes, or colliders. If the final non-endpoint node on an inducing path (let us denote it as 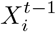) is not in the set of hidden nodes, then it must be a collider. If it is a collider, then it must be caused by the final node on the path (let us denote it as 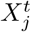), i.e., 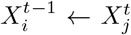. This is a cause backwards in time (since we have also assumed that there are no contemporaneous causes in the ts-DAG) which is not allowed, hence all final non-endpoint nodes on inducing paths must be in the set of hidden nodes, and so only variables with hidden nodes as a cause will receive additional edges in the ts-MAG. □

Because all hidden variables have themselves at the previous time step as a cause (Assumption 3), an inducing path can always be lengthened by adding additional auto-links. This means that the hidden-node-induced additional edges will therefore persist to arbitrarily large lags, so that for any choice of *τ*_max_, an edge into the boundary node will exist at all lags up to *τ*_max_ (see also [3]).

We will now state the lemmas of Proposition 1 characterizing the types of additional edges received by boundary nodes in the ts-MAG and in which cases they occur.

#### Lemma 1.

*If a hidden node is cause of an observed variable*, 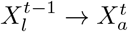 *in a ts-DAG, then there exist additional auto-link edges* 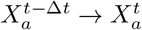 *in the ts-MAG for all* 1 *<Δt < τ*_*max*_.

*Proof of Lemma 1*. To guarantee the existence of an auto-link edge 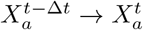 for all 1 *<Δt <τ*_*max*_, we must find an inducing path between 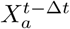 and 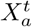for all 1 *<Δt < τ*_*max*_. If 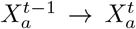 then 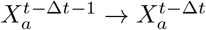for all Δ *t >* 1 due to the repeating structure of the network (and for all other edges).

An inducing path is then

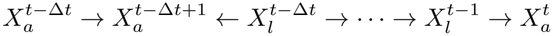

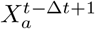 is a collider on the path and is an ancestor of the endpoint 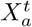. All other non-endpoint nodes on the path are hidden nodes of the variable *X*_*l*_. □

Lemma 1 implies that boundary subnetwork nodes that have a hidden variable as a cause will acquire delayed auto-link edges in the ts-MAG for all 1 *<Δt <τ*_*max*_.

#### Example 1.

*The example of Fig. 1a shows a ts-DAG with two variables, X*_*l*_ *and X*_*a*_. *X*_*l*_ *causes X*_*a*_ *with a lag of one. The ts-MAG of Fig. 1b is the resulting network when X*_*l*_ *is hidden, as a result of Lemma 1*.

**Figure 1:**
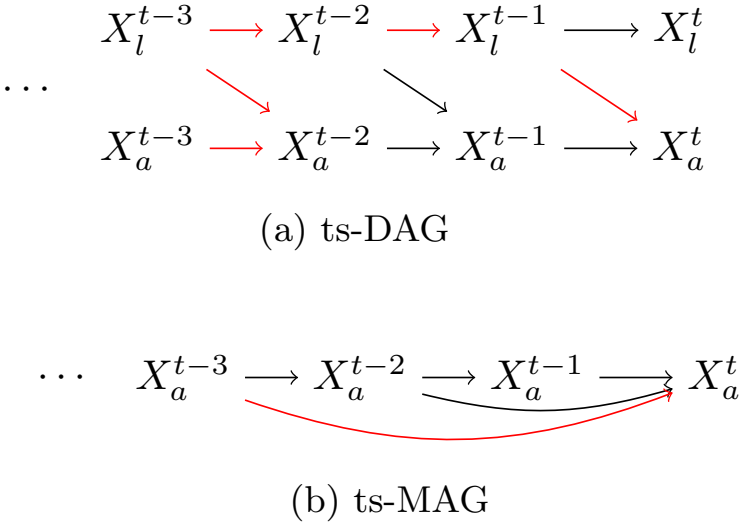
Hidden node as cause of a boundary node. Example inducing path for 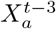 to 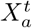in the ts-DAG shown in red, and corresponding edge 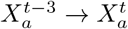 in the ts-MAG shown also in red.

#### Lemma 2.

*If a hidden node is an intermediate between two observed variables*, 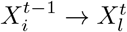 *and* 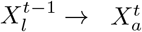 *in a ts-DAG, then there exist additional cross-link edges* 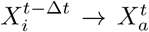 *in the ts-MAG for all* 1 *<Δt <τ*_*max*_.

*Proof of Lemma 2*. To guarantee the existence of a cross-link edge 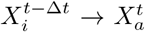 in the ts-MAG for all 1 *<Δt <τ*_*max*_., we must find an inducing path between 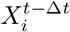 and 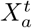for all 1 *<Δt < τ*_*max*_. If 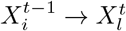then 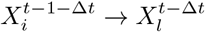 for all 1 *<Δt < τ*_*max*_.due to the repeating structure of the network.

An inducing path is then

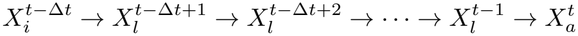

i.e. all non-endpoint nodes on the path are hidden nodes of the variable *X*_*l*_. □

Lemma 2 implies that when a hidden variable acts as an intermediate between two observed nodes, the boundary node will acquire a delayed cross-link edge in the ts-MAG for all 1 *<Δt < τ*_*max*_.

#### Example 2.

*The example of Fig*. 2a *shows a ts-DAG with three variables, X*_*i*_, *X*_*l*_ *and X*_*a*_. *X*_*i*_ *causes X*_*l*_, *which then causes X*_*a*_ *each with a lag of 1. The ts-MAG of Fig. 2b is the resulting network when X*_*l*_ *is hidden, as a result of Lemma 2*.

**Figure 2:**
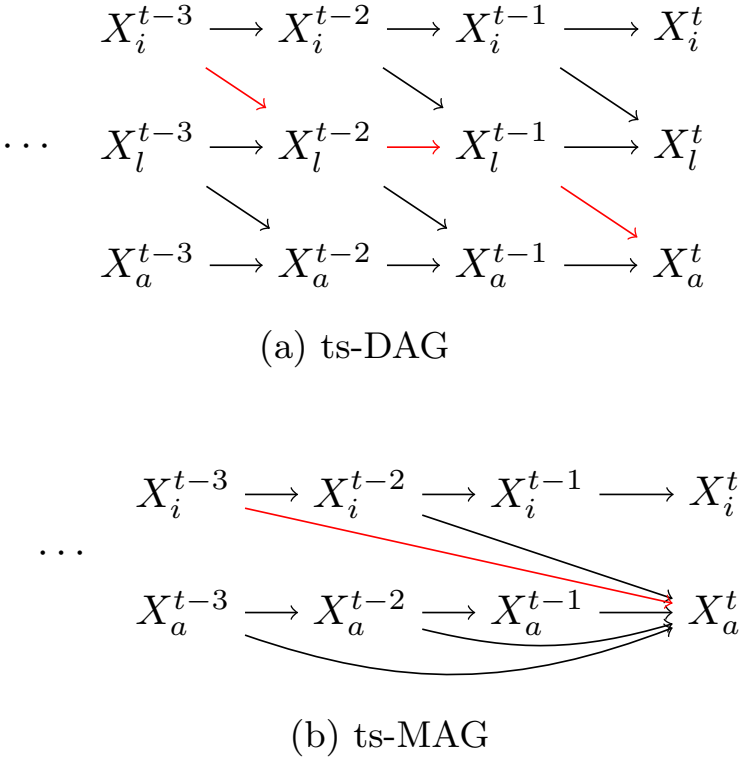
Hidden node as an intermediate between observed variables. Example inducing path for 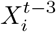 to 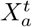in the ts-DAG shown in red, and corresponding edge 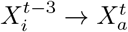 in the ts-MAG shown also in red.

#### Lemma 3.

*If a hidden node is an autocorrelated confounder, i*.*e*., 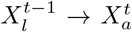 *and* 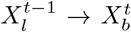, *then each boundary node caused by X*_*l*_ *will receive additional edges from each boundary node in the ts-MAG at all lags up to τ*_*max*_. *That is*, 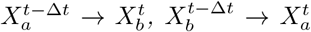 *and* 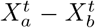 *for all* 1 *<Δt < τ*_*max*_.*and* 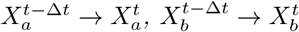*for all* 1 *<Δt < τ*_*max*_.

*Proof*. To guarantee the existence of edges 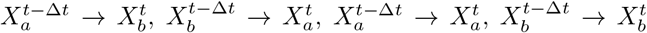 and 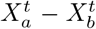 the ts-MAG for all 1 *<Δt < τ*_*max*_, we must find inducing paths between each of the corresponding nodes for all 1 *<Δt < τ*_*max*_.

For the edge 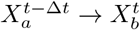, we must find an inducing path between 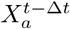 and 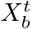. An inducing path is then

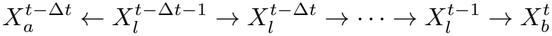

i.e., all non-endpoint nodes on the path are hidden nodes *X*_*l*_.

For the edge 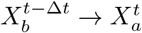 the inducing path is as above, but with the indices *a* and *b* exchanged.

For the auto-link edge 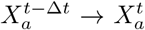, the inducing path is as above, but with the indices *b* replaced with *a*.

For the auto-link edge 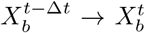 the inducing path is as above, but with the indices *a* replaced with *b*.

The inducing path between 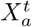and 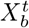is simply

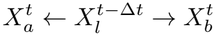

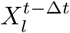 is a hidden node, hence 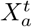and 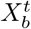are adjacent in the ts-MAG. □

Lemma 3 implies that when a hidden variable acts as a confounder between two observed nodes, each of the boundary nodes will acquire cross-link edges from each of the other boundary nodes in the ts-MAG for all *Δt < τ*_*max*_.

#### Example 3.

*The example of Fig*. 3a *shows a ts-DAG with three variables, X*_*l*_, *X*_*a*_ *and X*_*b*_. *X*_*l*_ *causes both X*_*a*_ *and X*_*b*_ *with a lag of 1. The ts-MAG of Fig. 3b is the resulting network when X*_*l*_ *is hidden, as a result of Lemma 3*.

**Figure 3:**
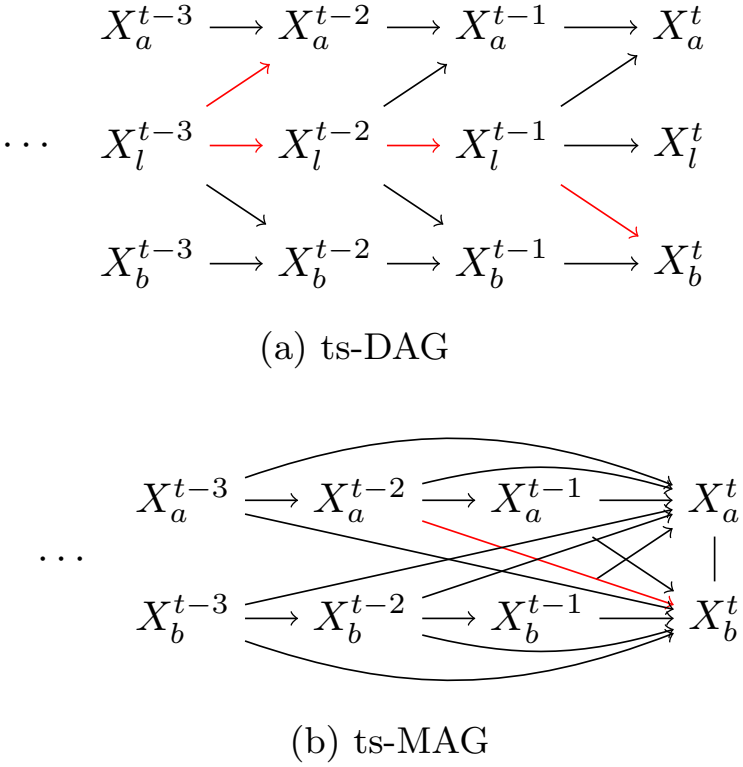
Hidden node as a confounder of two observed variables. Example inducing path for 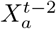 to 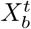 in the ts-DAG shown in red, and corresponding edge 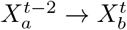 in the ts-MAG shown also in red.

#### Lemma 4.

*If a boundary node has any other cause or causes along with the hidden node*, 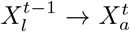 *and* 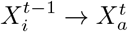, *it will receive lagged edges from these other causes*, 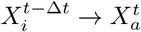 *for all* 1 *<Δt < τ*_*max*_.

*Proof*. To guarantee the existence of a cross-link edge 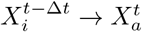 in the ts-MAG for all 1 *<Δt < τ*_*max*_,

we must find an inducing path between 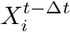 and 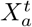. This inducing path is then

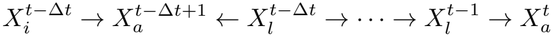

i.e., 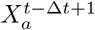 is a collider on the path and ancestral to 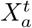, while all other non-endpoint nodes on the path are hidden nodes of *X*_*l*_. □

Lemma 4 implies that when a boundary species has multiple causes (including nodes that are not hidden), the boundary node will acquire cross-link edges from each of these causes in the ts-MAG for all 1 *<Δt τ*_max_.

#### Example 4.

*The example of Fig. 4a shows a ts-DAG with three variables, X*_*i*_, *X*_*l*_ *and X*_*a*_. *Both X*_*l*_ *and X*_*i*_ *cause X*_*a*_ *with a lag of 1. The ts-MAG of Fig. 4b is the resulting network when X*_*l*_ *is hidden, as a result of Lemma 4*.

**Figure 4:**
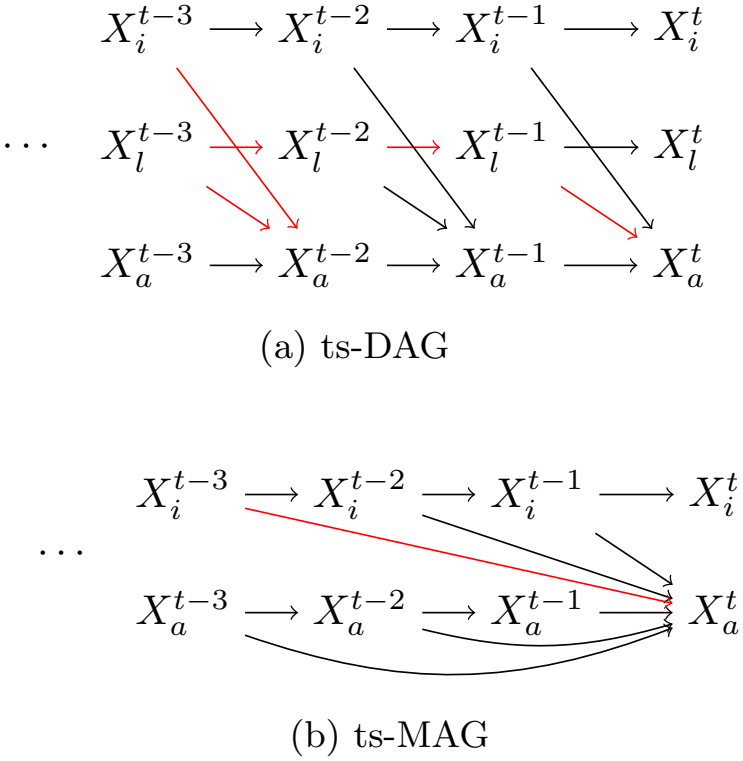
Boundary node with multiple causes. Example inducing path for 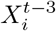 to 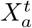in the ts-DAG shown in red, and corresponding edge 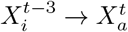 in the ts-MAG shown also in red.

#### Full adjacency matrix of the ts-MAG

Since we have shown that additional edges induced in the ts-MAG can be inferred purely from the network structure of the ts-DAG, and that the network structure is described by the matrix ***K***, it follows then that the adjacency matrix ***A*** of the ts-DAG specified by ***K*** can be used to infer the additional edges in the ts-MAG. ***Ã*** is the block adjacency matrix of the subnetwork-reduced time-unfolded skeleton causal network. Each block is of size *N* ^s^ × *N* ^s^ with the block ***Ã***_*t,t−*Δ*t*_ representing links from time *t−* Δ *t* to *t* and each element can be computed as follows.

Edges to internal subnetwork nodes, *X*^*i*^ (i.e., subnetwork nodes which do not interact with hidden nodes), from time *t* − 1 to *t* are given by the edges in the existing ts-DAG

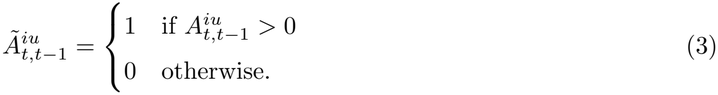

While at all other times there are no edges into internal subnetwork nodes

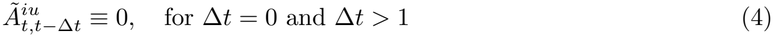

where *X*_*u*_ may be either an internal subnetwork node, or a boundary node.

Edges to boundary subnetwork nodes, *X*^*a*^, at the same time step (*t* to *t*) are due to confounding hidden nodes are given by

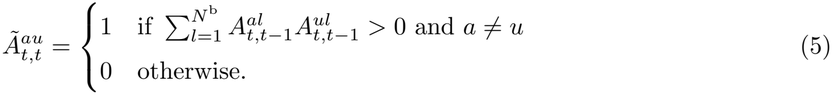

Edges into boundary subnetwork nodes from *t* −1 to *t* are given by existing subnetwork edges and by any effects from confounders

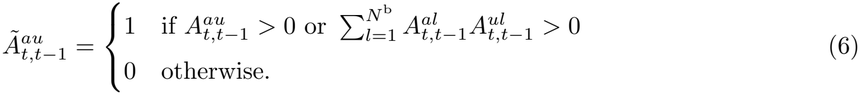

Edges into boundary subnetwork nodes from other lags *t* − Δ *t* into *t* for 1 *<Δt < τ*_*max*_ come from hidden nodes as intermediates, hidden nodes as confounders, and from any other causes of boundary nodes

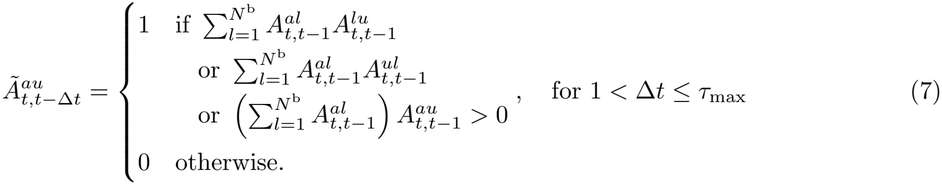

We can also write expressions for the blocks of ***Ã*** in terms of the matrix ***K*** (as in the main text), since ***A***_*t,t−*1_ is non-zero wherever ***K*** is non-zero

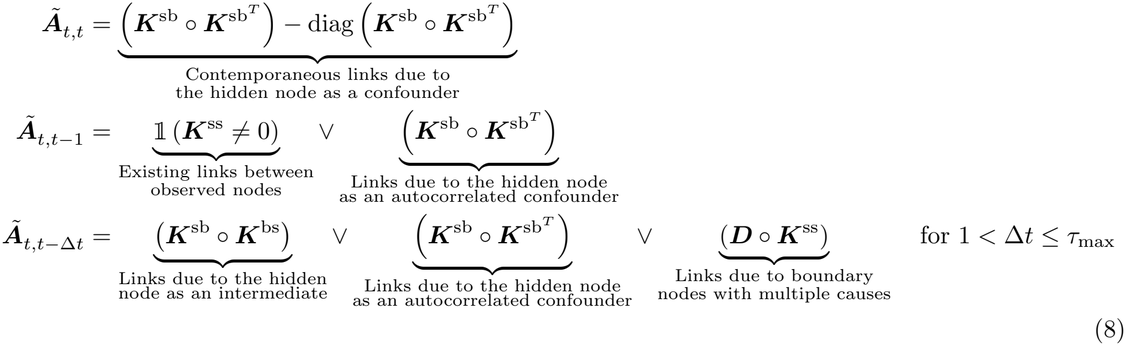

where 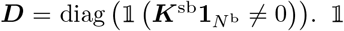 . is an indicator function, 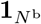is the vector of *N* ^b^ ones so that ***D*** is an *N* ^s^ × *N* ^s^ diagonal matrix with ***D***^*uu*^ = 1 if and only if *X*_*u*_ is a boundary node. ⋁ is the logical ‘or’ operation and the boolean matrix operation ° is defined as (***U*** ° ***V***)_*ij*_ = ^ _*k*_ ((*U*_*ik*_ ≠ 0) ^ (*V*_*kj*_ ≠ 0)) with ^ being the logical ‘and’.

